# Postprandial profiling of the duodenal microbiome reveals the impact of food structure and association with luminal metabolite and gut hormone responses

**DOI:** 10.64898/2026.05.06.723166

**Authors:** Frederick J. Warren, Katerina Petropoulou, Hannah C. Harris, Cecia Barbas-Bernardos, Melpomeni Kasapi, Antonia Garcia, Elaine Holmes, Claire Domoney, Julien Wist, Isabel Garcia-Perez, Gary Frost

## Abstract

The human duodenum harbours a complex, dynamic microbial community that is challenging to study due to inaccessibility, particularly postprandially when nutrient-rich chyme and fluctuating metabolites create unique microbial niches. We used naso-duodenal intubation to longitudinally sample duodenal luminal contents following pea-based meals of differing food structure, alongside parallel blood collection. Shotgun metagenomic sequencing, comprehensive metabolomic profiling and gut hormone measurements were combined to explore microbe-metabolite-hormone interactions. Food structure significantly affected postprandial bacterial composition, with saccharolytic oral taxa increasing after meals with intact structure. Alpha diversity was influenced by structure type (P = 0.025), with whole pea seeds promoting greater diversity than pea flour. Network analysis revealed complex interactions between the duodenal microbiome, luminal metabolites and gut hormones, with most microbial associations linked to glucose-dependent insulinotropic polypeptide (GIP) rather than glucagon-like peptide-1 (GLP-1). Metabolic profiling showed meal-dependent changes in amino acid metabolism, including shifts in D/L amino acid ratios over time consistent with microbial metabolism. The duodenal microbiome showed close phylogenetic relationships with the oral microbiome, with composition influenced by food structuring and swallowing. These findings reveal dynamic microbe-metabolite interplay in the human duodenum during digestion and its relationship to gut hormone responses.

## Introduction

The human gut microbiome is a diverse and complex microbial ecosystem, which plays an important role in food digestion, as well as maintaining homeostasis and health of the host.^1^ The majority of gut microbiome studies have focussed on the colon as this is the most densely populated part of the gastrointestinal tract, and also the easiest to access for sampling through stool samples, however stool samples do not capture host-microbe interactions *in situ*.^2^ The small intestinal microbiome has been understudied due to the challenges of accessing and sampling the lumen of the small intestine for analysis, and to date the majority of studies have utilised 16S amplicon sequencing, rather than shotgun metagenomic approaches.^2,3^ Previous studies have revealed that there are distinct segment-specific microbial communities within the small intestine, with a core small intestinal microbiome found in all compartments consisting of *Streptococcus, Veillonella, Fusobacterium, Prevotella* and *Haemophillus* and a duodenal segment specific community containing *Neisseria, Granulicatella, Rothia* and *Gemella*.^3,4^ However, the lack of species resolved data and functional insight has limited our understanding of the microbial community in the small intestine. The gastrointestinal tract can be considered as a continuum, with flow from the oral cavity to the ileum and beyond onto the colon. Many of the bacterial genera typical of the oral cavity such as *Streptococcus, Haemophilus, Rothia, Neisseria* and *Veillonella* are also found in the duodenum, suggesting a potential transmission route, although this remains to be proven.^5,6^ The small intestinal microbiome has also been shown to be highly saccharolytic and recent sequencing studies using capsule-based sampling have revealed that there is a wide diversity of Carbohydrate Active EnZymes (CAZymes) represented in the genomes of bacteria in the small intestine.^3,7^ There is abundant substrate available for saccharolytic bacteria in the small intestine as food passes through and the products of digestion are released, such as small sugars. However, the metabolic activity and carbohydrate degrading roles of the microbiome in the small intestine remain to be characterised in detail.^3^

It is well established that the duodenum plays a key role in nutrient sensing and gut hormone secretion. It contains enteroendocrine K and L cells, which release the incretin hormones GIP and GLP-1, respectively, in response to the absorption of nutrients such as glucose and fatty acids from the intestinal lumen.^8^ The microbiome can impact this signalling, both directly, through the production of short chain fatty acids (SCFA) which are detected by enteroendocrine cells,^9^ and indirectly, through modulating the metabolic environment of the gut.^10,11^ The effect of the microbiome on gut hormone signalling is well characterised in the colon,^9^ but not so in the small intestine.

In a previous study,^12^ we carried out an intervention trial using two different pea genotypes, one with a naturally occurring mutation termed the *rugosus* (*rr*, wrinkled-seeded) genotype, and the other a near-isogenic control (*RR*, round-seeded) genotype. The *r* mutation is in the Starch Branching Enzyme 1 (*SBE1*) and leads to very high amylose starch, with altered granule morphology and thermal properties.^13^As a consequence, the *rr* mutant pea seeds have significantly lower starch digestibility, resulting in a lower post-prandial glycaemic response, lower insulinemic response, and also lower GLP-1 and GIP in the study participants.^12^ As part of the current intervention trial investigating starch-microbiome-metabolome interactions in the small intestine, a naso-duodenal intubation was carried out allowing direct sampling of the contents from the duodenum. Participants consumed a meal containing either whole pea seeds from the two different genotypes or the corresponding milled pea flours. Across a series of experiments investigating the impact of the *r* mutation and food structural properties, we observed that the *rr* pea led to a significantly reduced release of glucose into the duodenum compared with the control genotype.^12^ In the present study we present an analysis using species-resolved shotgun metagenomics of the microbiome composition of the duodenum. We explored the dynamics of the duodenal microbiome during the consumption of pea-based meals and examined how the food structure, either in whole peas or pea flour, influences microbial composition. Our analyses revealed associations between microbial taxa, metabolites, functional profiles, and gut hormone responses.

## Results

### Composition and dynamic community shifts of the duodenal gut microbiome

In the fasted state, the duodenal microbiome of the study participants was dominated by Firmicutes from the order Lactobacillales, with significant contributions from Proteobacteria, Bacteroidetes and Actinobacteria (Figure 1A and Supplementary Figure S1A). At the species level, the duodenal microbiome was dominated by taxa predominantly associated with the oral microbiome, including *Rothia mucilaginosa*, *Haemophilus parainfluenzae*, *Prevotella jejuni*, *Gemella sanguinis*, and several *Streptococcus* species (*S. infantis, S. mitis, S. salivarius*). The fasted microbial community, sampled on four independent occasions for each participant, was relatively stable within individuals but showed significant inter-individual variation (PERMANOVA, F = 1.8059, p = 0.001, Figure 1B), demonstrating individual-specific duodenal microbiome signatures with some intra-individual variability between sampling occasions.

**Figure 1.**
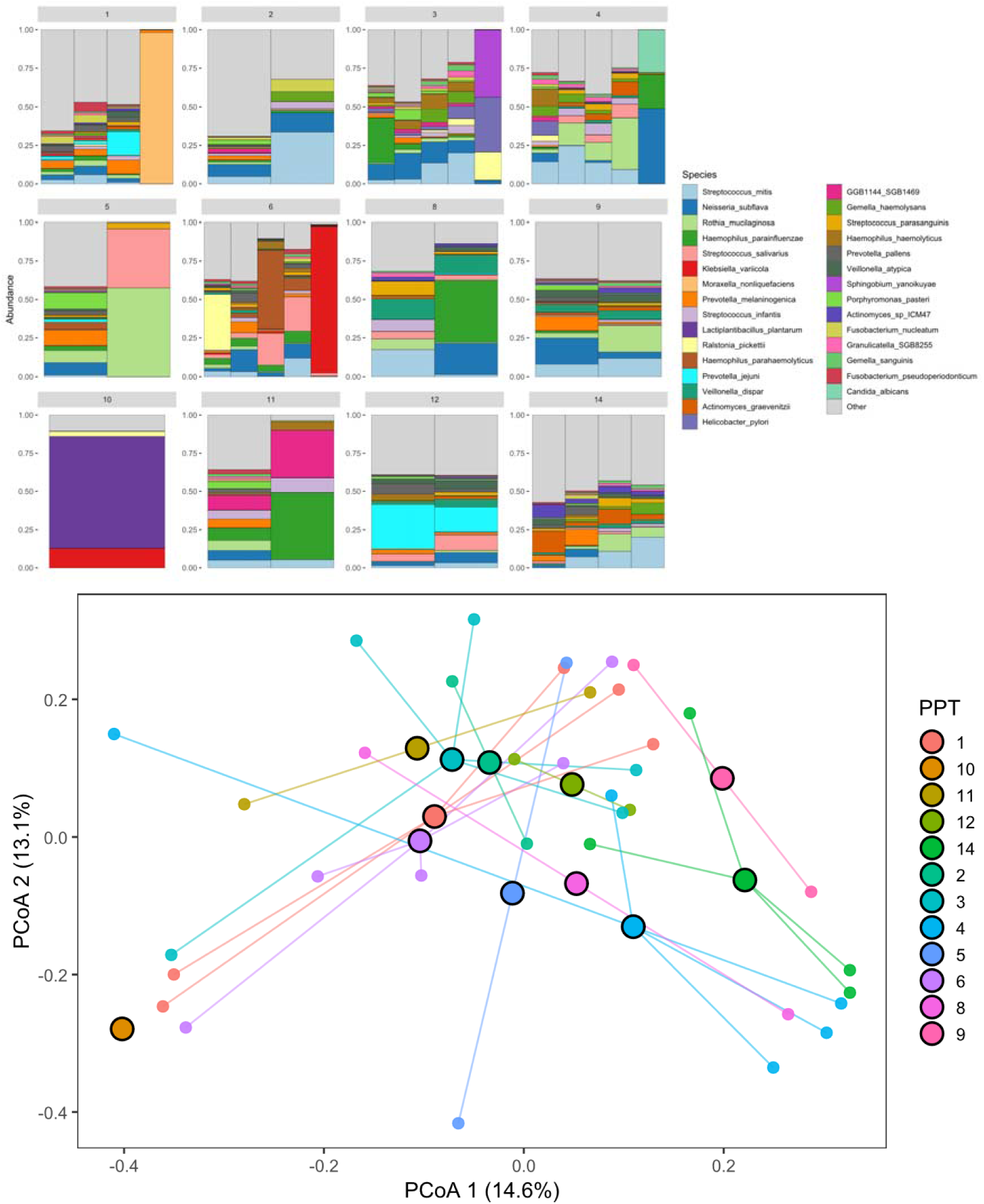
Baseline (fasted) microbiome composition of the duodenal lumen. **A.** Microbiome profiles of each participant for samples taken at baseline. The relative abundance of the top 30 most abundant species is shown. **B.** PCoA plot (Bray-Curtis dissimilarity) showing the community composition of the duodenal microbiome of the participants at baseline. The data are coloured by participant number, with participant centroids highlighted by black circles. Note that for some donors baseline data was not available from all meals.

Following meal consumption, the duodenal microbiome remained broadly stable in terms of overall diversity. Alpha diversity (Shannon index) was significantly influenced by food type (F = 5.08, p = 0.025), with whole peas promoting greater diversity than flour, but not influenced by time following meal consumption or genotype alone (Figure 2). However, species-level changes were observed in a subset of taxa, with significant increases in *Streptococcus cristatus* (F = 22.99, p = <0.001) and *Gemella sanguinis* (F = 22.63, p = <0.001) over time (Figure 3 and 4). The passage of pea material from the stomach into the duodenum could be tracked through reads mapping to the Pisum sativum genome, which peak between 30 and 60 minutes post-consumption (Figure S1B), with higher proportions observed from pea flour samples compared to intact, whole pea seeds. This corresponds to the presence of the compounds trigonelline and stachyose, which are unique undigestible markers of peas,^14^ and which also peak in the duodenal between 30 and 60 minutes postprandially (Figure S7).

**Figure 2.**
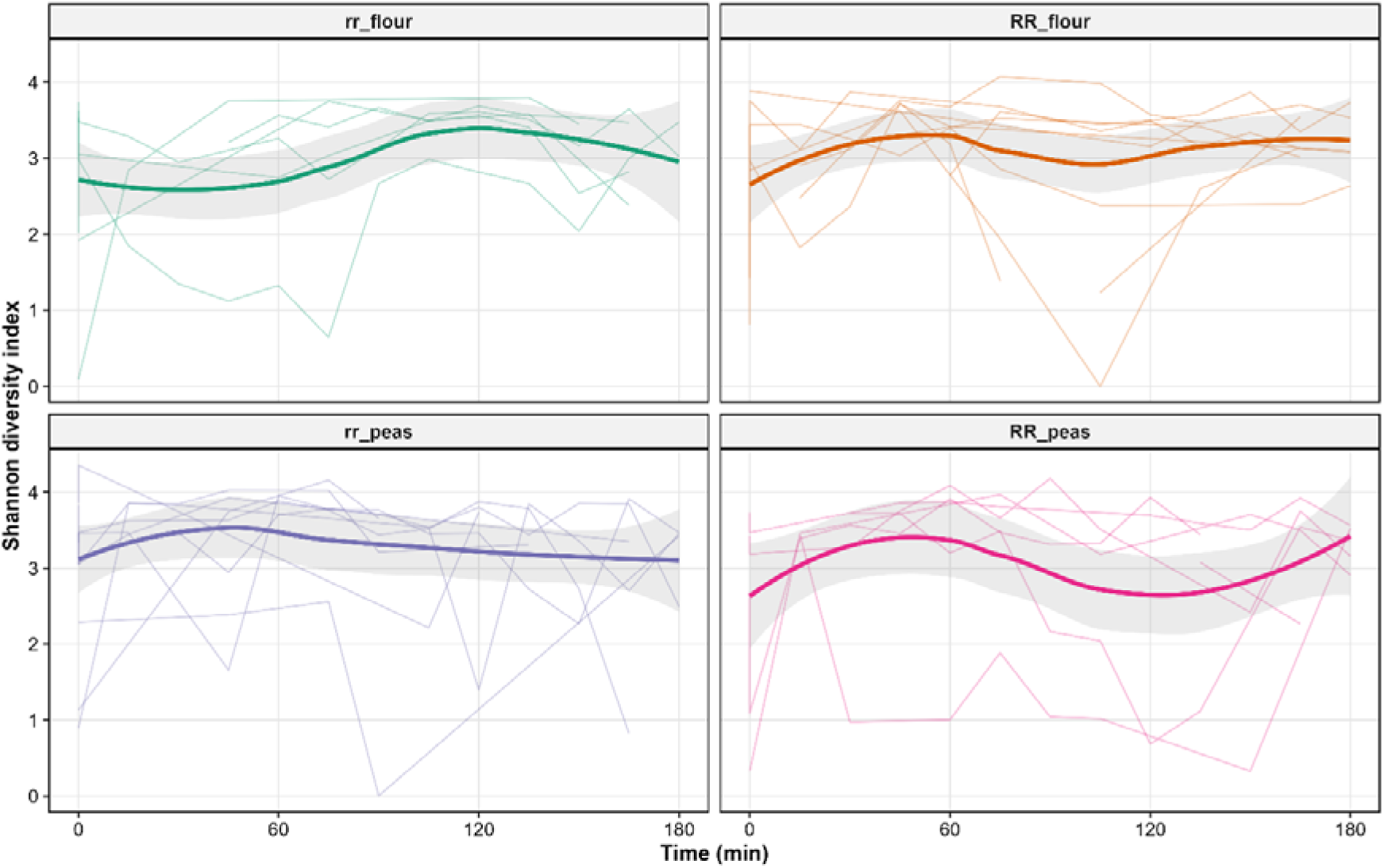
Alpha diversity changes over time by meal type. Shannon diversity over time in the duodenal microbiome following consumption of four different meals. Individual participant trajectories are shown as thin lines (n = 14 participants per meal), with LOESS-smoothed mean trajectories ± standard error shown as thick lines and shaded regions. Time 0 represents baseline before meal consumption.

**Figure 3.**
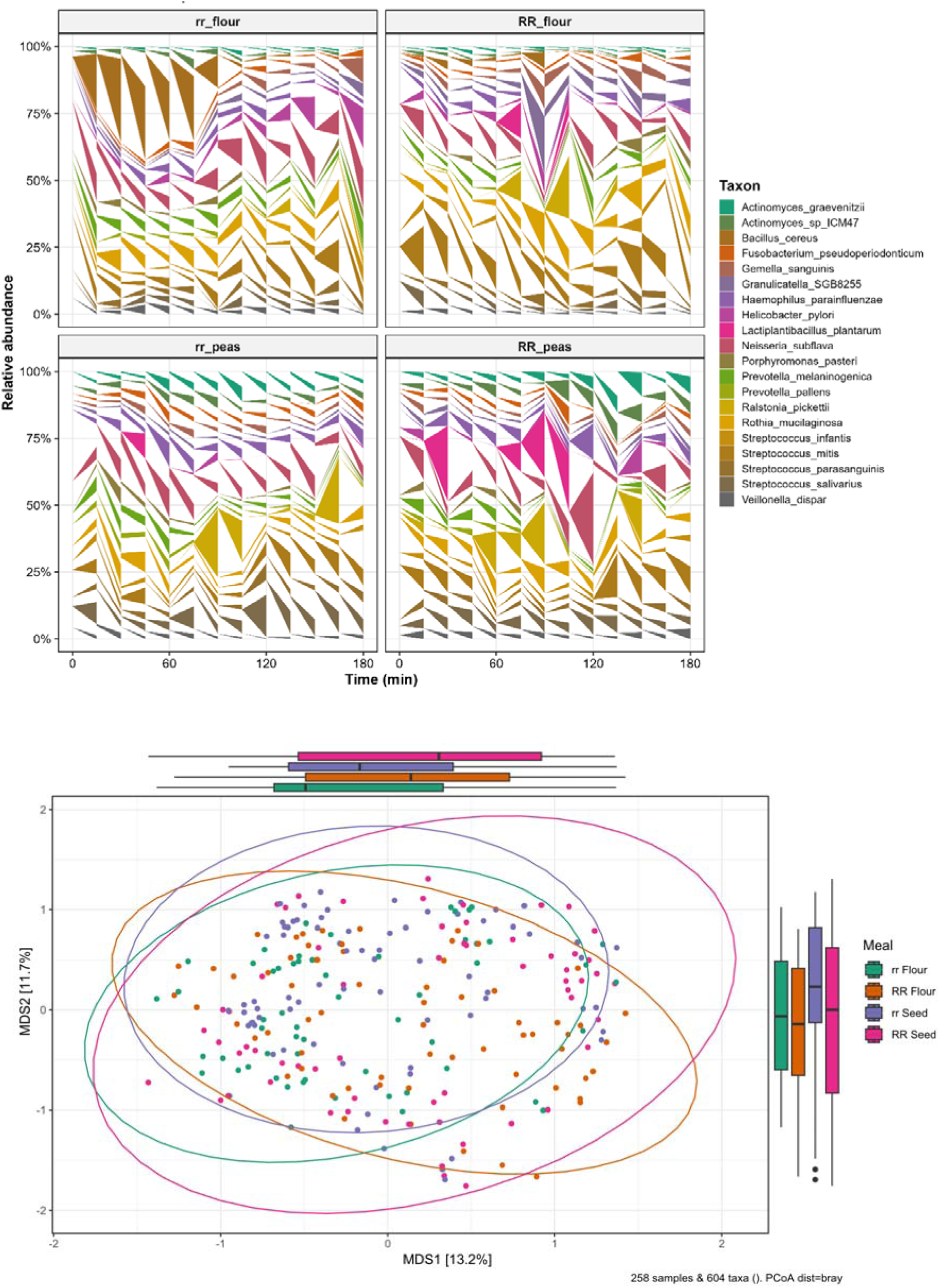
Compositional and beta-diversity changes in the duodenal microbiome in response to pea meals. **A.** Area plot showing the relative abundance of the top 20 species in the duodenal microbiome, and their changes in abundance over time following consumption of each of the meals. Values are averaged across the participants. **B.** PCoA plot showing the microbial community composition (Bray-Curtis dissimilarity) following consumption of each of the pea meals. The colour indicates the meal type.

**Figure 4.**
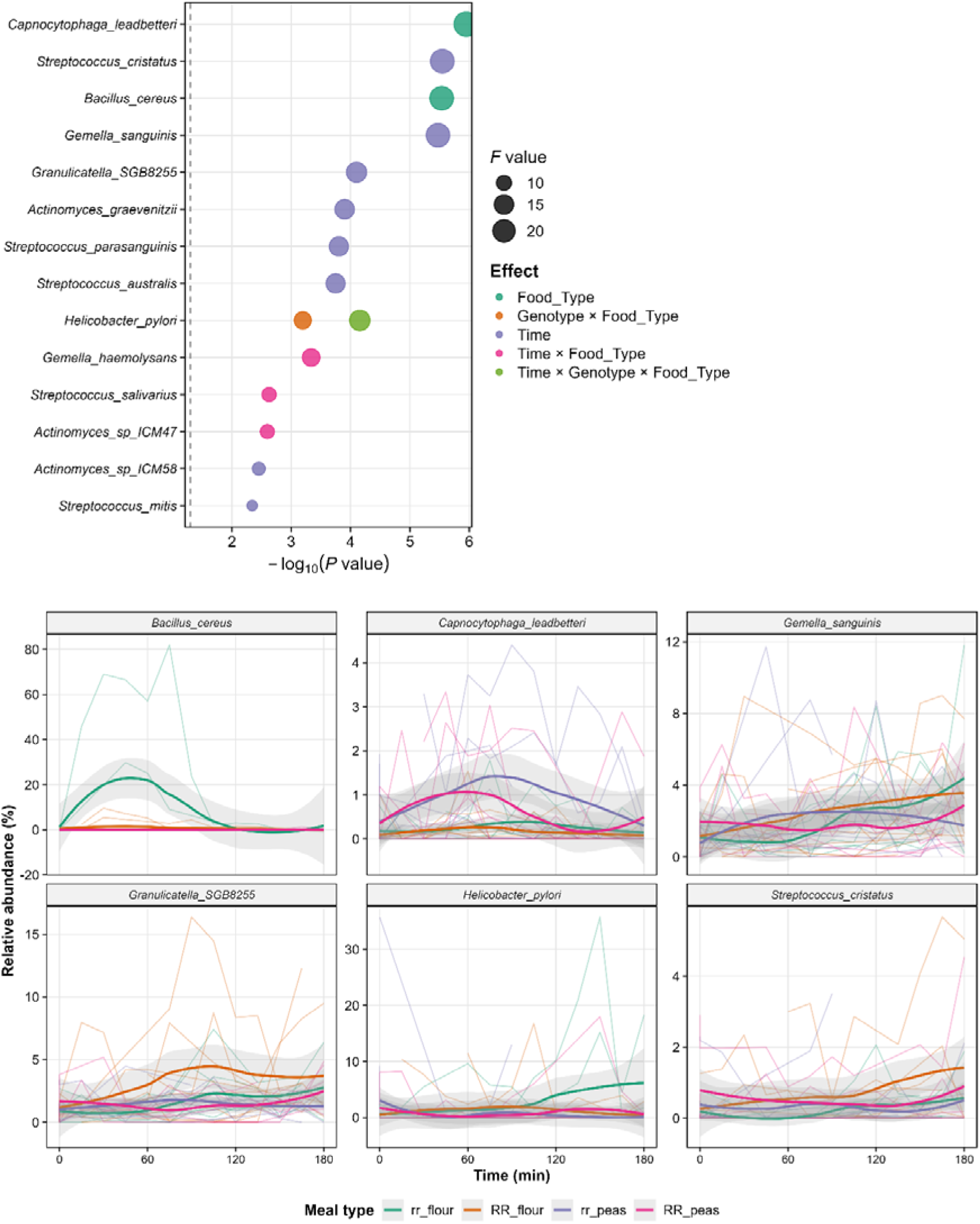
Differential abundance analysis reveals meal- and time-dependent bacterial responses. Identification and characterization of significantly differentially abundant taxa. (A) Dot plot showing the 15 most significantly differentially abundant taxa (ranked top to bottom by P-value). Horizontal position indicates significance (-log□□ P-value), with the vertical dashed line marking P = 0.05. Point size represents effect size (F-value from ANOVA), and color indicates the type of effect (green = time, orange = genotype, purple = food type, magenta = genotype × food type interaction, etc.). Multiple points per taxon indicate responses to multiple experimental factors. (B) Abundance trajectories for the six most significantly affected taxa across the four meal types. Individual participant trajectories (thin lines) and LOESS-smoothed means ± standard error (thick lines and shaded regions) are shown.

### Duodenal microbiome composition is influenced by food structure

Two food structural variables were explored: peas provided as whole seeds (highly structured) or milled flour (less structured), and two pea genotypes (*rr* mutant with high amylose content versus *RR* wildtype). Community composition differed significantly based on both physical form and genotype (PERMANOVA: R² = 0.365, F = 9.97, p = 0.001). Physical form had a larger effect (F = 6.85, p = 0.001) than genotype (F = 3.11, p = 0.002) on microbial community composition (Figure 3B, Supplementary Figures S3-S4). In the fasted state (prior to consumption of the peas and following an overnight fast), no differences were observed between interventions. Following meal consumption, divergence between food structures became apparent within 30-60 minutes post-prandially (Figures S2 and S3).

Differential abundance analysis identified numerous species responding to experimental factors (Figure 4A, Supplementary Figure S4). More species changed in response to physical form than genotype. Notably, *Capnocytophaga leadbetteri* (F = 24.93, p = <0.001) and *Bacillus cereus* (F = 22.91, p = <0.001) were strongly associated with food type. The strong association of *B.cereus* with only one food type may indicate that this is a contribution from the food microbiome as this is common food contaminant. Multiple *Streptococcus* species showed time-dependent changes, with *S. cristatus, S. parasanguinis, and S. australis* all significantly increasing post-prandially.

### Metabolic profiling reveals dynamic changes in the duodenal luminal environment

Metabolic phenotyping of duodenal aspirates by ¹H NMR spectroscopy identified a diverse array of metabolites including amino acids, organic acids, sugars, and other small molecules. The metabolite profiles showed meal (food and genotype)-dependent and time-dependent variation, reflecting the dynamic nature of the postprandial duodenal environment (Figure S5).

Amino acids represented a major class of metabolites detected in the duodenal lumen, with both proteinogenic and non-proteinogenic forms present. Amino acid concentrations peaked later than sugars in the sampling period, from 120 to 180 minutes, and there was a trend towards higher amino acid release from the intact peas, although this was non-significant. Sugars, including glucose and maltose, were released during the earlier stages of digestion peaking between 30 and 60 minutes, with a significant difference in maltose depending on food structure, with higher concentrations observed for the flours compared to the intact peas (Figure S5).

Organic acids including lactate, acetate, propionate and formate were also detected in duodenal aspirate, although at low concentrations. Succinate and formate showed significant differences depending on food structure, higher in the flour compared to the intact peas. The presence of short-chain fatty acids in the duodenal lumen, though at lower concentrations than typically observed in the colon, suggests active microbial fermentation even in this proximal region of the gastrointestinal tract. These metabolites may contribute to the local metabolic environment and potentially influence enteroendocrine cell signalling.

### Network analysis reveals complex microbe-metabolite-hormone interactions in the duodenum

To comprehensively characterize the relationships between duodenal microbiota, luminal metabolites, and gut hormones, we performed integrated network analysis using Spearman rank correlations on False Discovery Rate (FDR)-corrected significant associations (q < 0.05, Figure 5). The network comprised 40 bacterial species, 17 metabolites, and 2 gut hormones (GIP and GLP-1), connected by 240 significant correlations. The network revealed distinct communities of interacting features, with clear separation between GIP-associated and GLP-1-associated microbial taxa, and notably more extensive microbial associations with GIP (14 connections) than GLP-1 (10 connections). This pattern aligns with the duodenal location being the primary site of GIP secretion, with GLP-1 predominantly secreted more distally in the jejunum and ileum.

**Figure 5.**
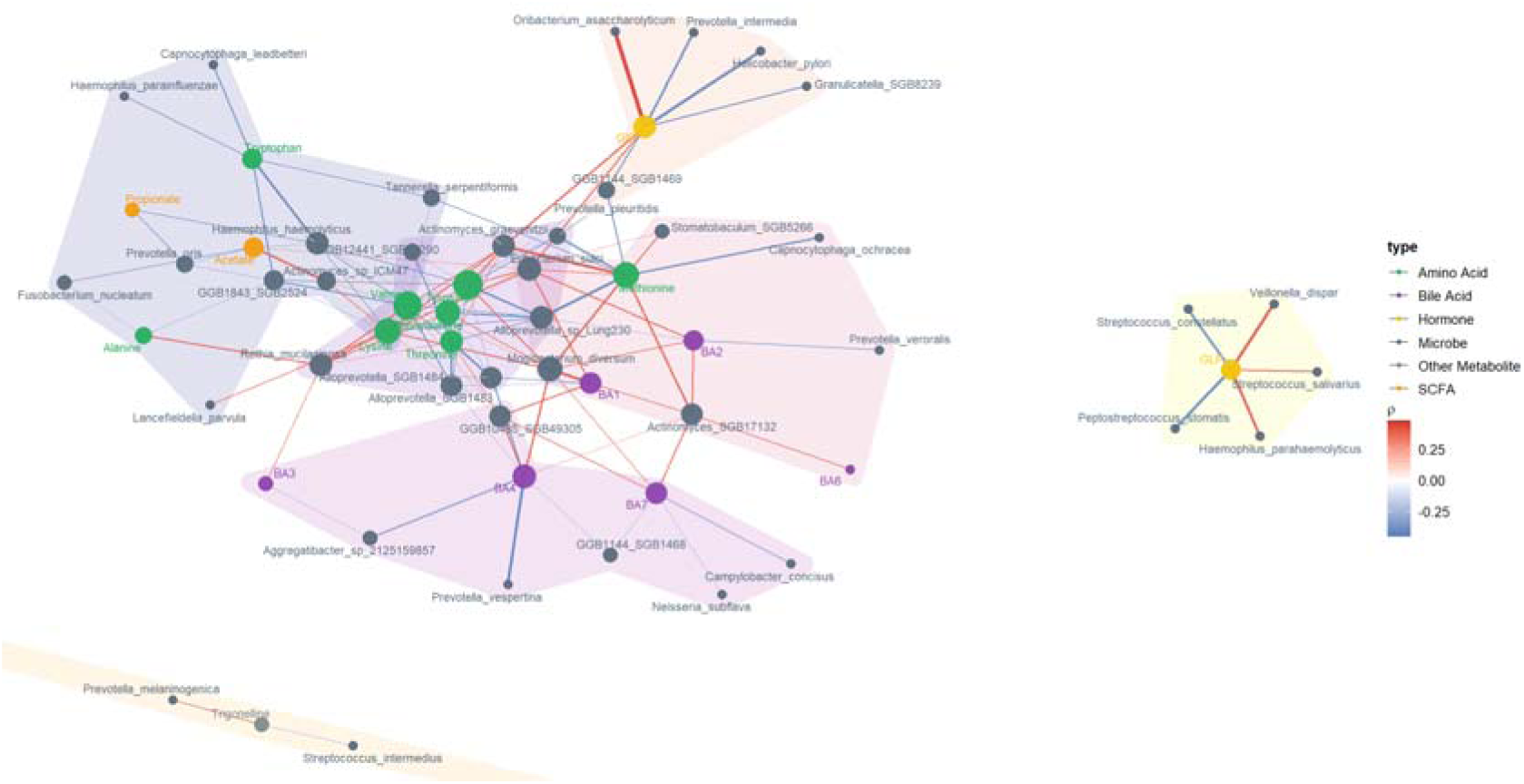
Network analysis of hormone-metabolite-microbe interactions. Spearman rank correlated network showing interactions in the lumen of the duodenum between z-scored metabolite concentrations (blue), gut hormone levels (magenta) and centred log-ratio (CLR) normalised microbial abundances (green). Edges represent significant correlations (fdr corrected *p* < 0.05), with edge colour indicating the direction and strength of the correlation. The size of the node indicates relative abundance(bacteria) or concentration (metabolites and hormones). Shaded areas delineate network communities detected using the Louvain modularity algorithm.

The network was organized into seven distinct communities detected using the Louvain modularity algorithm, revealing clusters of co-occurring and functionally related features. The two smallest communities were hormone-centric, centered around GIP and GLP-1 respectively. GIP formed a tightly connected module with five bacterial species, *Oribacterium asaccharolyticum, Eubacterium sulci,* and *Actinomyces graevenitzii* showing positive correlations with GIP, while *Helicobacter pylori, Prevotella intermedia, Granulicatella SGB8239*, and *GGB1144 SGB1469* showed negative correlations. GLP-1 formed a separate community with *Streptococcus salivarius*, *Veillonella dispar*, and *Haemophilus parahaemolyticus* (all positive correlations), and *Peptostreptococcus stomatis* and *Streptococcus constellatus* (negative correlations). The clear separation of GIP and GLP-1 into distinct network communities, with no overlap in their associated bacterial species, reflects the anatomical and functional distinction between these incretin hormones in the duodenum.^8^

Several bacterial species emerged as key network hubs based on their degree (number of connections) and betweenness centrality. The most connected hubs were *Mogibacterium diversum* and *Eubacterium sulci*, both of which showed extensive correlations with metabolites and positive associations with GIP. Other prominent hubs included *Haemophilus haemolyticus*, which clustered with amino acids including lysine, tryptophan, alanine, and organic acids (acetate, propionate). *Rothia mucilaginosa* and *Actinomyces graevenitzii* formed a hub in a community characterized by branched-chain and aromatic amino acids (valine, tyrosine, phenylalanine) and threonine, representing a distinct metabolic niche focused on protein degradation products. These findings demonstrate that the duodenal microbiome contains specific taxa that act as metabolic hubs, with extensive connections to luminal metabolites and in some cases to gut hormone signaling.

The remaining metabolite-microbe communities revealed functional clustering patterns, including a community clustering bile acid-related metabolites (BA3, BA4, BA7) with *Neisseria*, *Campylobacter*, and *Aggregatibacter* species. The modularity of the network, with distinct communities centred around different metabolite classes and hormone systems, indicates that the duodenal microbiome exhibits functional specialization with different bacterial consortia associated with specific metabolic processes and host signaling pathways.

### D/L amino acid ratios reveal bacterial amino acid metabolism in the duodenum

A particularly striking finding from the metabolomic analysis was the quantification of D-amino acids alongside their L-enantiomers in the duodenal lumen. D-amino acids are typically produced by bacterial amino acid racemases and are rare in eukaryotic systems, making them useful markers of bacterial metabolic activity.^15^ Although several animal studies have successfully quantified free D-AAs concentrations within the gut lumen, research in humans and especially in the duodenal lumen remains scarce.^16^ We calculated D/L ratios for several amino acids and observed significant temporal changes following meal consumption (Figure 6A).

**Figure 6.**
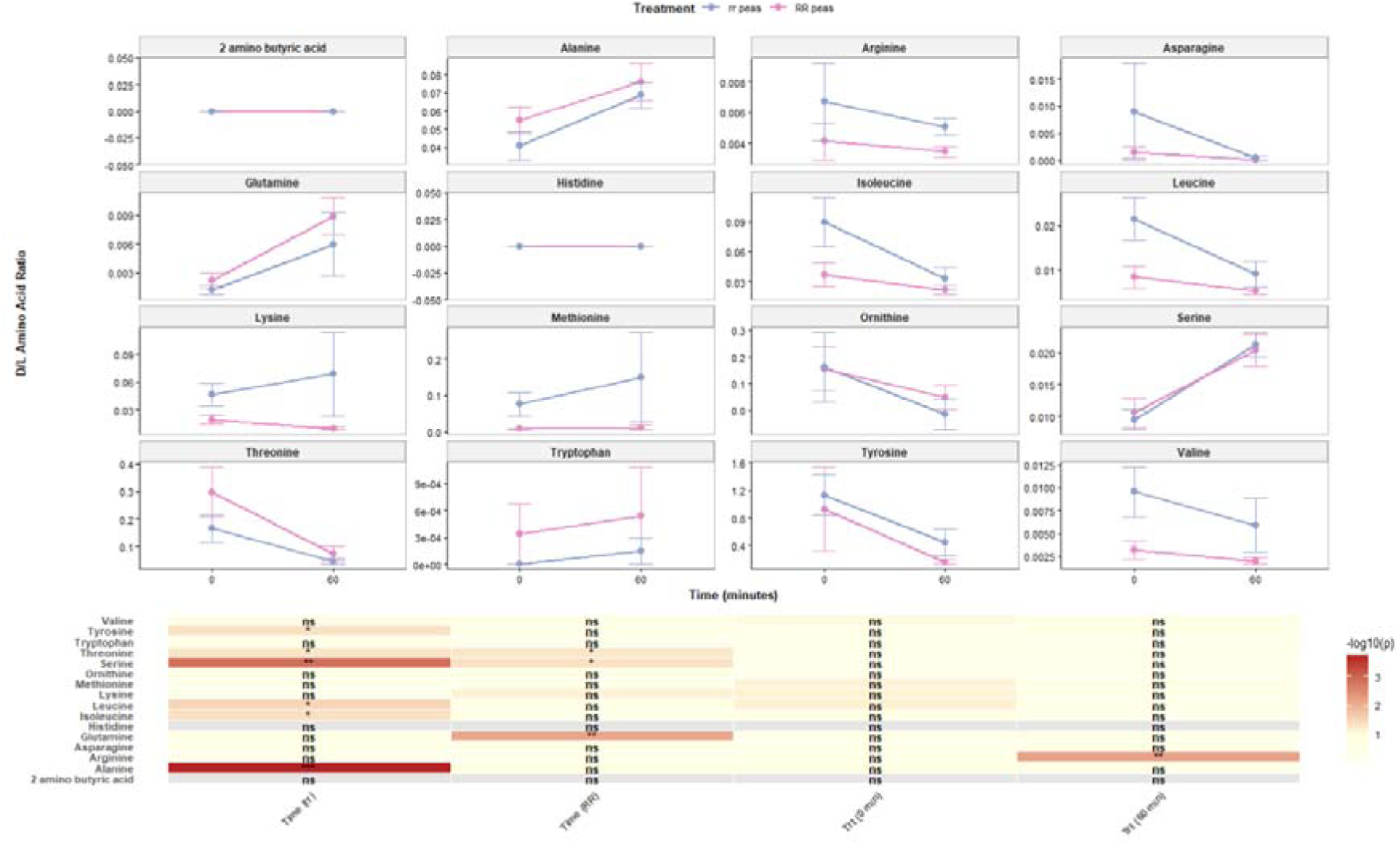
The interconversion of L and D amino acids in the lumen of the duodenum. (A) Changes over time (from baseline to 60 minutes) in the D/L amino acid ratio in the lumen of the duodenum, following consumption of either the rr or RR pea meals. (B) Differences in DL amino acid ratios over time for each meal, determined by paired t-tests, and between meals at each timepoint, determined by independent t-tests.

For multiple amino acids including alanine, serine, aspartate, and glutamate, the D/L ratio increased significantly from baseline to 60 minutes post-consumption (paired t-tests, p < 0.05). This temporal pattern suggests ongoing bacterial racemization of L-amino acids released during meal digestion. Interestingly, the magnitude and kinetics of D/L ratio changes differed between the *rr* and *RR* pea meals (Figure 6B). The *rr* peas, which have more resistant starch and maintain greater structural integrity, showed more gradual increases in D/L ratios, potentially reflecting slower substrate release and different patterns of bacterial colonization or activity on the food particles.

The production of D-amino acids in the duodenum has several potential functional implications. D-amino acids can act as signaling molecules in bacterial biofilms, modulating biofilm assembly and dispersal. They may also be incorporated into bacterial peptidoglycan, particularly in stationary phase cells.^15^ Furthermore, D-amino acids have been shown to interact with host signalling pathways, including effects on neutrophil chemotaxis and potentially on interactions with taste receptors expressed on enteroendocrine cells.^17,18^ The correlation between D-amino acid levels and specific bacterial taxa in the network analysis, particularly *Streptococcus* and *Veillonella* species known to possess amino acid racemases (Figure S6), provides evidence for active bacterial amino acid metabolism in the human duodenum and suggests another layer of microbe-host biochemical interaction.

### CAZyme profiles reflect carbohydrate availability and microbial adaptation to food structure

Given the starch-rich nature of the pea meals and the associations between microbiome composition, glucose release, and gut hormones, we analyzed the Carbohydrate Active Enzyme (CAZyme) profiles of the duodenal microbiome using functional metagenomic annotation (Supplementary Figure S7). The CAZyme analysis revealed significant associations between specific enzyme families and experimental variables, with the greatest number of associations observed for physical meal form (whole peas versus flour), consistent with the microbiome composition results.

Two CAZyme families showed particularly strong and functionally relevant patterns. Glycoside Hydrolase family 4 (GH4) showed positive associations with both luminal glucose concentrations and consumption of whole peas. GH4 enzymes exhibit diverse catalytic activities, but the most common function is α-glucosidase activity, which degrades maltose and maltooligosaccharides to glucose. Some GH4 enzymes are also involved in sucrose metabolism and other carbohydrate transformations.^19^ The association of GH4 with whole peas may reflect the higher availability of partially digested starch granules and oligosaccharides as substrates for bacterial degradation.

Carbohydrate Binding Module family 41 (CBM41) showed strong positive associations with GIP hormone levels and consumption of whole peas. CBM41 domains are characterized by their ability to bind various forms of starch, including amylose, amylopectin, and their degradation products. These modules are typically found appended to starch-degrading enzymes, enhancing their ability to target and degrade starch substrates.^20^ The association between CBM41 abundance and GIP secretion (Supplementary Figure S7) is intriguing and may reflect the role of starch-binding organisms in modulating the rate and location of carbohydrate digestion, thereby influencing glucose release and incretin hormone responses.

Taxonomic analysis of the distribution of these CAZyme families revealed distinct patterns (Supplementary Figure S7). CBM41 was predominantly encoded by *S. sanguinis* and *S. salivarius*, two oral streptococci that are abundant in the duodenum and known for their ability to metabolize complex carbohydrates. In contrast, GH4 showed broader taxonomic distribution across multiple genera including *Leptotrichia, Klebsiella, Enterobacter*, and *Citrobacter*. Both CAZyme families showed marked increases in abundance following consumption of whole peas compared to pea flour. This suggests that the physical structure of food rapidly influences not only which bacteria are present but also their functional capabilities for carbohydrate metabolism.

## Discussion

The duodenal microbiome has mainly been studied in the context of disease, due to the challenge of accessing the duodenum for sampling.^2,21^ Relatively few studies have attempted to sample the duodenal microbiota in healthy participants. Here, we have presented the duodenal microbiota of 12 healthy participants with species level resolution, alongside comprehensive profiling of the duodenal metabolome and network analysis, revealing complex microbe-metabolite-hormone interactions.

Previous studies, primarily relying on 16S sequencing, have revealed that the duodenal microbiota is quite distinct from the microbial community composition found in stool, and is dominated by the genera *Streptococcus, Actinomyces, Veillonella, Gemella* and *Rothia*, with significant variability between individuals.^22^ The findings of this study generally confirm these findings, extending to species resolution. Within the genus *Streptococcus,* the species *salivarius, mitis, infantis* and *parasanguinis* were the most abundant, along with *R. mucilaginosa, G. sanguinis* and *haemolysans*, *A. graevenitzii* and *V. dispar, atypica* and *rogosae*. We also found high abundance of the *Haemophilus* species *parainfluenzae* and *parahaemolyticus*. There was a broad overlap between the species which were observed at high abundance in the duodenum and those which are typical of the mucosal regions (tongue dorsum and buccal mucosa) of the mouth, suggesting there may be a link between these communities.^23^

Previous studies of the duodenal microbiome have been limited through their sampling approaches to only studying the fasted state. In the present study we achieved intensive sampling of the duodenal microbiome during ingestion of structurally diverse pea-based meals from two different genetic backgrounds and two contrasting food structures.^12^ We demonstrated that the degree of structuring of the pea (whether it is presented as a whole pea or as flour), as well as the genotype effects, influenced the microbiome composition of the duodenum. This builds on our previous findings that the pea structure and genotype could influence glucose release in the duodenum and gut hormone responses.^12^ A key advance in this study is the integration of metagenomic, metabolomic, and hormone data through network analysis, which revealed distinct communities of interacting features and highlighted the central role of GIP in duodenal microbe-host communication.^24^ The associations observed between duodenal microbiome and hormone responses are suggestive, particularly as the majority of associations were with GIP, which is primarily released in the duodenum, rather than GLP-1 which is released further down the GI tract in the jejunum and ileum.^8,25^ The network analysis identified key microbial taxa such as *A. graevenitzii*, *E. sulci*, and *O. asaccharolyticum* (positively correlated with GIP) and *H. pylori* and *P. intermedia* (negatively correlated with GIP) as potential mediators of these interactions. However, with the currently available data we cannot state if these associations are causative, i.e. if the microbiome of the duodenum is directly influencing hormone release, or if the microbiome changes are associated with food structure or duodenal metabolic profiles, which lead to differing hormone responses.

This study represents the first integrated metagenomic and metabolomic analysis of the human duodenal microbiome in a healthy cohort during consumption of a meal. The duodenal microbial community is dominated by *Streptococcus, Actinomyces, Veillonella, Gemella* and *Rothia*, and the species present have a strong taxonomic overlap with the oral mucosal microbiome, with significant interindividual variability. We demonstrate that the duodenal microbiome is impacted by the ingestion of food and that the physical structure of the food shapes the microbial community in the duodenum. Through network analysis, we reveal complex tripartite interactions among the microbiome, luminal metabolites, and gut hormones. The metabolomic data provide insights into the dynamic chemical environment of the duodenum and potential microbial contributions through amino acid metabolism. Finally, we demonstrate that there are associations between GIP release from the duodenum and duodenal microbiome composition, suggesting a possible role for microbial activity in nutrient sensing in the gut. These findings highlight the duodenum as a critical site for microbe-host interactions that may influence postprandial metabolism and hormone responses.

## Methods

### Human clinical trial

The food materials used were as follows: wild-type pea seeds (BC1/19RR line), used as a control group; mutant pea seeds (BC1/19*rr* line), used as a treatment group; wild-type pea flour (BC1/19RR line), used as a control group; mutant pea flour (BC1/19*rr* line), used as a treatment group.

The human intervention trials were conducted and samples collected as described in Petropoulou et al.^12^ Briefly, twelve healthy volunteers (men and women aged 18–65 years, with a body mass index of 18.5–29.9 kg/m^2^) took part in a randomised, controlled, double-blind, four-way crossover study, attending the Clinical Research Facility for four consecutive days (three nights), with each test meal administered in random order on a separate study day. Naso-gastric and naso-duodenal tubes were placed by a clinician using the CORPAK system (MedSystems, Halyard) and remained in place for the duration of the four-day visit. On each study day, after an overnight fast, two fasting baseline samples (blood and gastric/duodenal aspirate) were collected at –10 and 0 min. At 0 min, volunteers consumed 50 g dry weight of RR or rr pea seeds or pea flour as the sole test meal (i.e. without an accompanying mixed meal). Postprandial blood, gastric and duodenal aspirate samples were then collected at 15, 30, 60, 90, 120 and 180 min. Volunteers were instructed to consume a standardised meal the evening before each study visit and to avoid caffeine, alcohol and strenuous exercise for 24 h prior to each procedure. All study visits took place at the National Institute for Health Research (NIHR)/Wellcome Trust Imperial Clinical Research Facility, Hammersmith Hospital, London, UK and were conducted between May 2015 and December 2017. All studies were approved by the South East Coast Surrey Research Ethics Committee (15/L0/0184) and carried out in accordance with the Declaration of Helsinki. Luminal glucose, serum GLP-1 and serum GIP were measured as described in Petropoulou et al.^12^

### Chiral analysis and quantification

Duodenal samples were thawed and centrifuged (13,000 × *g* for 5 min at 4 °C). For the extraction of polar compounds, a 100 µL aliquot of the supernatant was gently mixed with 1.0 mL of methanol and centrifuged (13,000 × *g* for 10 min at 4 °C). Subsequently, 100 µL of the resulting supernatant was mixed with 750 µL of cold acetonitrile and 25 µL of internal standard (IS, 25 µmol/L), vortexed, and centrifuged again (20,000g for 5 min at 4 °C). Then, 150 µL of the supernatant was transferred to an Eppendorf tube for the derivatization step. To concentrate the sample, the supernatant was evaporated to dryness using a SpeedVac concentrator for 1 h at 45°C. The residue was resuspended in 50 µL of water and 35 µL of 0.15 mol/L sodium tetraborate (pH 9), and the solution was mixed for 10 min. For derivatization, 50 µL of (S)-NIFE (12.5 mg/mL in acetonitrile) was added, mixed, and allowed to react for 10 min. The reaction was stopped by adding 10 µL of cold 4 mol/L HCl and 355 µL of water. Samples were kept on ice for 10 min and then centrifuged (13,000 × *g* for 10 min at 4 °C). Finally, the supernatant was transferred to vials. Quality control (QC) samples were prepared by pooling equal volumes of the supernatant from each sample.

The analysis was performed using an Acquity UPLC system coupled to a Xevo Triple Quadrupole mass spectrometer (Waters Corp.). A 5 µL aliquot of derivatized sample was injected onto a reversed-phase column maintained at 60 °C (Acquity BEH C18, 100 mm × 2.1 mm, 1.7 µm), equipped with a pre-column (Acquity UPLC BEH C18 VanGuard, 2.1 × 5 mm, 1.7 µm). The mobile phases consisted of (A) 10 mM ammonium bicarbonate in water (pH 9.5) and (B) acetonitrile, delivered at a flow rate of 0.6 mL/min with a total run time of 27 min. The gradient elution profile was as follows (time [min] / %B): 0/4, 5/4, 9/10, 11/11.5, 16/28, 19/30, 21/35, 22.5/90, 23.5/90, 25/4, and 27/4. Data were acquired in multiple reaction monitoring mode using an electrospray ionization source operated in positive mode. The capillary voltage was set to 2.5 kV, the cone voltage to 40 V, and the collision voltage and collision energy were optimized for each amino acid. The desolvation gas flow was 900 L/h and the cone gas flow was 40 L/h, with a desolvation temperature of 500 °C. Data acquisition was carried out using Waters MassLynx v4.1 software.

Quantification was performed using the stable isotope dilution method^26^ (Area_AA/Area_AA*), considering the corresponding D- or L-labeled amino acid (AA*). When the labeled D-enantiomer was not available, the corresponding L-labeled standard was used. For compounds lacking an internal standard (His, Pro, Orn, Cys, and Tyr), calibration curves were prepared for each enantiomer and used for quantification. Calibration curves were acquired at the beginning and end of the analytical sequence, as well as every two days for compounds without labeled standards. QC samples were also analyzed at the beginning of the sequence and after every ten injections to assess system stability and reproducibility.

Absolute quantification of D- and L-amino acids (µmol/L) was based on their relative peak areas with respect to the corresponding labeled internal standard and the known concentration of the internal standard, according to the equation: C (µmol/L) = (Area_AA / Area_IS) × C_IS × (10 / 4), where the dilution factor (DF = 4) accounts for mixing with the internal standard. When labeled standards were available as racemic mixtures (DL-AA*), the concentration of the L-enantiomer (L-AA*) in the working solution was determined by triplicate analysis against a fresh solution of the corresponding unlabeled L-amino acid, using the equation: C_L-AA* = (Area_L-AA* × C_L-AA) / Area_L-AA. The concentration of the D-enantiomer (D-AA*) was then calculated as the difference between the total labeled concentration and the L-enantiomer concentration (C_D-AA* = C_T-AA* − C_L-AA*). This approach was also applied when only the L-labeled standard (L-AA*) was available.

For amino acids lacking D-labeled standards, relative response factors (RRFs) between D- and L-enantiomers were determined by analyzing triplicate solutions containing 200 ng/mL of each pure enantiomer. The RRF was calculated as the ratio of the peak areas (Area_D-AA / Area_L-AA). In these cases, D-amino acid concentrations were calculated using the equation: C (µmol/L) = (Area_AA / Area_L-AA*) × C_L-AA* × (10 / (4 × RRF)).

### 1H NMR metabolic profiling

As previously described by Petropoulou et al. ^12^, duodenal samples were centrifuged (15 min, 3000 g) and metabolites extracted using a modified Folch method. Briefly, chloroform/methanol (2:1) was added to samples, followed by water to induce phase separation. After centrifugation (20 min, 3000 g, 0°C), the aqueous phase was collected, dried, and stored at −80°C.

For analysis, samples were reconstituted in water, sonicated, and mixed with phosphate buffer (pH 7.4, 80% D₂O) containing 3-(Trimethylsilyl)propionic-2,2,3,3 acid sodium salt (TSP) as an internal standard prior to transfer into NMR tubes. Duodenal samples were prepared similarly with minor adjustments in volumes and buffer concentration. Quality control samples were generated by pooling aliquots of each sample.

¹H-NMR spectroscopy was performed at 300 K on a 600 MHz Bruker spectrometer using a standard one-dimensional pulse sequence incorporating the first increment oof the NOE pulse sequence to achieve water suppression. Spectra were acquired with 32 scans (following 4 dummy scans), a spectral width of 20,000 Hz, and 64 K data points. Free induction decays were processed with 0.3 Hz line broadening prior to Fourier transformation. Spectra were manually corrected, referenced to TSP at δ 0.0, and regions corresponding to water and TSP were excluded prior to modelling the data. Data were processed in MATLAB and normalized using median fold change normalization with the median spectrum as reference.

### Metabolite identification and Quantification

Following established protocols,^27^ statistical spectroscopic methods, including statistical total correlation spectroscopy (STOCSY) and subset optimization by reference matching (STORM), were applied for signal identification. Metabolite assignments were confirmed using internal references and external databases, including the Human Metabolome Database (HMDB) (HMDB; http://hmdb.ca/) and the Biological Magnetic Resonance Data Bank (BMRB) (https://bmrb.io/).

A total of 24 metabolites were quantified using in-house routines developed in both JavaScript and R. NMR spectra were processed in accordance with the Bruker IVDr standard procedures,^28^ and no additional processing was deemed necessary. For quantification, metabolites of interest were first grouped into spectral regions of interest (SROIs). Within each SROI, a set of signals was defined with attributes describing their characteristic patterns, including chemical shift, linewidth, multiplicity, and scalar coupling; all detected patterns were included, even those corresponding to unknown signals. These parameters were then used as initial inputs for a gradient descent optimization algorithm (Levenberg–Marquardt) to minimize the difference between the experimental spectra and the fitted patterns. The resulting models were evaluated using a comprehensive heuristic that incorporated both optimization error metrics and measures of correlation among integrals from signals known to originate from the same compound. Models with lower scores were subsequently examined through visual inspection using a suite of interactive web-based tools developed in JavaScript.^29^ Finally, the calculated areas under the curve were converted into concentrations using the ERETIC internal standard and calibration procedure included in the Bruker IVDr protocol.

### DNA extraction and metagenomic sequencing

DNA extraction was carried out per the manufacturer’s instructions with the MP Bio fast DNA spin kit for soil (MP Biomedical, Solon, USA). Duodenal aspirates were transferred to Lysing Matrix E tubes where 980 µL of sodium phosphate buffer and 120□µL of MT buffer were added. The samples were homogenised in the FastPrep24 bead-beating instrument (MP Biomedicals, Solon USA) for 3□min (3 runs of 60□s each, with 5□min rest on ice in between). Afterwards, samples were centrifuged at 14,000□×□g□for 15□min, and the sample supernatant was transferred into clean 2□mL microcentrifuge tubes with 250□µL of protein precipitate solution. The tubes were mixed by shaking by hand 10 times before centrifugation at 14,000□×□g□for a further 10□min. The supernatant was transferred to 5□mL tubes with a resuspended binding matrix (1□mL of silica slurry that binds DNA from lysates), and the tubes were inverted for 2□min allowing the DNA to bind. The tubes were placed in a rack and allowed to settle for 1□h. Then, 500□µL of the supernatant was removed, and the binding matrix was resuspended in the remaining supernatant. The washing steps involved the transfer of 700□µL of the mixture to a SPIN filter centrifuged at 1000□×□g□for 2□min to empty the catch tube and transfer the leftover mix. This process was repeated until all the mixture had been added to the SPIN filter. The SPIN filter was washed with 500□µL of ethanol-based wash solution SEWS-M. The samples were centrifuged at 14,000□×□g□for 5□min, and the catch tube was emptied. Then, samples were centrifuged a second time (Dry spin) at 14,000□×□g□for 5□min to remove the residual wash solution. The catch tube was replaced with a clean tube (1.5□mL LoBind Eppendorf®□tubes), and samples in the SPIN filter were air-dried for 10□min at room temperature and 5□min at 37□°C. Then, 65□µL of DNA extraction solution was added to the Binding Matrix and incubated at room temperature for 5□min before being centrifuged at 6600□×□g□for 2□min with lids open to bring eluted DNA into the tube. The SPIN Filter was discarded, and the eluted DNA was stored at −20□°C.

Genomic DNA was normalised to 5□ng/µL with elution buffer (10□mM Tris HCl). A miniaturised reaction was set up using the Nextera DNA Flex Library Prep Kit (Illumina, Cambridge, UK). 0.5□µL Tagmentation Buffer 1 (TB1) was mixed with 0.5□µL Bead-Linked Transposomes (BLT) and 4.0□µL PCR-grade water in a master mix and 5□µL was added to each well of a chilled 96-well plate. About 2□µL of normalised DNA (10□ng total) was pipette-mixed with each well of Tagmentation master mix and the plate heated to 55°C for 15□min in a PCR block. A PCR master mix was made up using 4□µL kapa2G buffer, 0.4□µL dNTP’s, 0.08□µL Polymerase and 4.52□µL PCR-grade water, from the Kap2G Robust PCR kit (Sigma-Aldrich, Gillingham, UK) and 9□µL added to each well in a 96-well plate. About 2□µL each of P7 and P5 of Nextera XT Index Kit v2 index primers (catalogue No. FC-131-2001 to 2004; Illumina, Cambridge, UK) were also added to each well. Finally, the 7□µL of Tagmentation mix was added and mixed. The PCR was run at 72□°C for 3□min, 95□°C for 1□min, 14 cycles of 95□°C for 10□s, 55□°C for 20□s and 72□°C for 3□min. Following the PCR reaction, the libraries from each sample were quantified using the methods described earlier and the high sensitivity Quant-iT dsDNA Assay Kit. Libraries were pooled following quantification in equal quantities. The final pool was double-SPRI size selected between 0.5 and 0.7X bead volumes using KAPA Pure Beads (Roche, Wilmington, US). The final pool was quantified on a Qubit 3.0 instrument and run on a D5000 ScreenTape (Agilent, Waldbronn, DE) using the Agilent Tapestation 4200 to calculate the final library pool molarity. qPCR was done on an Applied Biosystems StepOne Plus machine. Samples quantified were diluted 1 in 10,000. A PCR master mix was prepared using 10□µL KAPA SYBR FAST qPCR Master Mix (2X) (Sigma-Aldrich, Gillingham, UK), 0.4□µL ROX High, 0.4□µL 10□μM forward primer, 0.4□µL 10□μM reverse primer, 4□µL template DNA, 4.8□µL PCR-grade water. The PCR programme was: 95□°C for 3□min, 40 cycles of 95□°C for 10□s, 60□°C for 30□s. Standards were made from a 10□nM stock of Phix, diluted in PCR-grade water. The standard range was 20, 2, 0.2, 0.02, 0.002, 0.0002□pmol. The pooled library was then sent to Novogene (Cambridge, UK) for sequencing using an Illumina NovaSeq 6000 instrument, with sample names and index combinations used. Demultiplexed FASTQ’s were returned on a hard drive. An average sequencing depth of ∼10.1 GB per sample was achieved.

### Bioinformatic analysis

The raw FASTQ files were quality controlled using the package KneadData. First repetitive reads were removed using the package TRF and the adaptors and low-quality reads were removed with Trimmomatic.^30^ Then host contamination was removed by mapping reads using Bowtie2^31^ to the human reference genome hg37 and the human reference transcriptome hg38. In addition, a large number of reads from the *Pisum sativum* genome were identified in some samples, and these were removed with reference to the *Pisum sativum* v1a genome.^32^ The statistics for read counts before and after quality control are available in Supplementary Table S1.

Taxonomic profiles were obtained for each of the samples using the MetaPhlAn 4 package.^33^ Functional metagenomic profiles were obtained using the package HUMAnN 3.0 and CAZyme profiles were obtained by mapping the outputs to UniRef.^34,35^

### Statistical analysis and network analysis

All statistical analyses were performed in R (version 4.5.1). Alpha diversity metrics (Shannon index) were calculated from relative abundance data. The effects of time, pea genotype (RR vs rr), and food type (whole peas vs flour) on alpha diversity were assessed using linear mixed-effects (LME) models fitted with the lme4 and lmerTest packages, with the model specified as Shannon ∼ Time × Genotype × Food_Type + (1|Participant), where participant was included as a random intercept to account for the repeated-measures crossover design.

Beta diversity was assessed using Bray–Curtis dissimilarity computed from relative abundance profiles using the phyloseq package. Differences in community composition between meal types, genotype, and food type were tested using permutational multivariate analysis of variance (PERMANOVA; adonis2 function, vegan package) with 999 permutations and a seed of 123 for reproducibility. The full model was specified as Distance ∼ Genotype × Food_Type + Participant. Pairwise PERMANOVA comparisons between all four meal types were performed with Benjamini–Hochberg (BH) correction for multiple testing. Principal coordinates analysis (PCoA) was used for ordination and visualisation of community composition, with 95% confidence ellipses plotted using Student’s t-distribution. Time-stratified PERMANOVA analyses were conducted at each sampling timepoint to identify when compositional differences between groups emerged, with BH correction applied across all timepoints.

Differential abundance analysis was performed on the 50 most abundant taxa. Relative abundances were arcsine square-root transformed to stabilise variance and improve normality of proportional data. For each taxon, an LME model was fitted: Abundancetransformed ∼ Time × Genotype × Food_Type + (1|Participant). Type III analysis of variance was performed on each model using Satterthwaite’s method for denominator degrees of freedom (lmerTest package), and effects with P < 0.05 were considered significant.

Metabolite concentrations measured by ¹H NMR spectroscopy and hormone concentrations (18 metabolites, 6 bile acid species, and 2 gut hormones: GIP and GLP-1) were z-score normalised prior to integration with microbiome data. Between-group differences in metabolite concentrations at each timepoint were assessed using Kruskal–Wallis rank-sum tests.

To assess microbiome–metabolite–hormone associations, microbiome abundance data were centred log-ratio (CLR) transformed using the compositions package to account for the compositional nature of the data. Z-scored metabolite and hormone concentrations were combined with CLR-transformed microbial abundances, and pairwise Spearman rank correlations were computed using the Hmisc package (rcorr function). P-values were corrected for multiple testing using the Benjamini–Hochberg false discovery rate (FDR) method. Correlations were considered significant at FDR-adjusted q < 0.05 with an absolute Spearman ρ > 0.3.

Significant correlations meeting both thresholds were used to construct an undirected weighted network using the igraph package. Nodes represented microbial taxa, metabolites, or hormones, and edges represented significant correlations, with edge weights corresponding to Spearman ρ values. Node types were classified as microbe, short-chain fatty acid (SCFA), amino acid, bile acid, hormone, or other metabolite. Network topology was characterised by computing degree, betweenness centrality, and closeness centrality for each node. Community structure was identified using the Louvain modularity maximisation algorithm (cluster_louvain), with communities representing groups of nodes that are more densely connected to each other than to the rest of the network. The network was visualised using the ggraph package with a Fruchterman–Reingold force-directed layout, with community membership indicated by shaded convex hulls (ggforce package). Edge colour was mapped to correlation strength and direction using a diverging blue–white–red gradient representing negative to positive correlations, respectively.

All visualisations were produced using ggplot2, with additional packages patchwork, ggrepel, ggtext, and pheatmap. The following R packages were used for specific analyses: phyloseq (microbiome data handling), vegan (ecological statistics), lme4 and lmerTest (mixed-effects modelling), igraph and ggraph (network analysis), Hmisc (correlations), compositions (CLR transformation), and ggforce (network visualisation).

## Acknowledgments

We thank Martina Tashkova for her help during nasoduodenal tubes insertion and Mai Khatib and Rasha Alshaalan for their help with samples collection and gut hormone analysis. The clinical trial was conducted at the NIHR/Wellcome Trust Imperial Clinical Research Facility. The Division of Integrative Systems Medicine and Digestive Disease at Imperial College London receives financial support from the National Institute of Health Research (NIHR) Imperial Biomedical Research Centre (BRC) based at Imperial College Healthcare NHS Trust and Imperial College London, in line with the Gut Health research theme. This work was also supported by the Ministry of Science, Innovation and Universities of Spain PID2024-162248NB-I00 MICIU/AEI /10.13039/501100011033/ and by ERDF ‘A way of making Europ’ (A.G.). The authors thanks David Baker (Quadram Institute) for assistance with library preparation and sequencing. The authors gratefully acknowledge the support of the Biotechnology and Biological Sciences Research Council (BBSRC); this research was funded by the BBSRC Institute Strategic Programme Food Microbiome and Health BB/X011054/1 and its constituent projects BBS/E/QU/230001B and BB/X018857/1. CD was supported by BBSRC grant BBS/E/J/000PR9799.

## Contributions

F.J.W. conceived the study, carried out the bioinformatics analysis and analysed and interpreted the data and wrote the manuscript. K.P. helped conceive the study and led the human intervention trial. H.C.H. carried out DNA extractions and sequencing. M.K. assisted with bioinformatics analysis. J.W., E.H. and I.G.-P. carried out the metabolomic analysis. A.G, I.G.-P. and C.B.-B carried out the chiral analysis. G.F. and C.D. supervised and conceived the project.

## Data availability

Raw sequence data are deposited in NCBI SRA under project number PRJNA1425766.

