## Supplementary material for "Postprandial profiling of the duodenal microbiome reveals the impact of food structure and association with luminal metabolite and gut hormone responses": Supplemenatary Figures

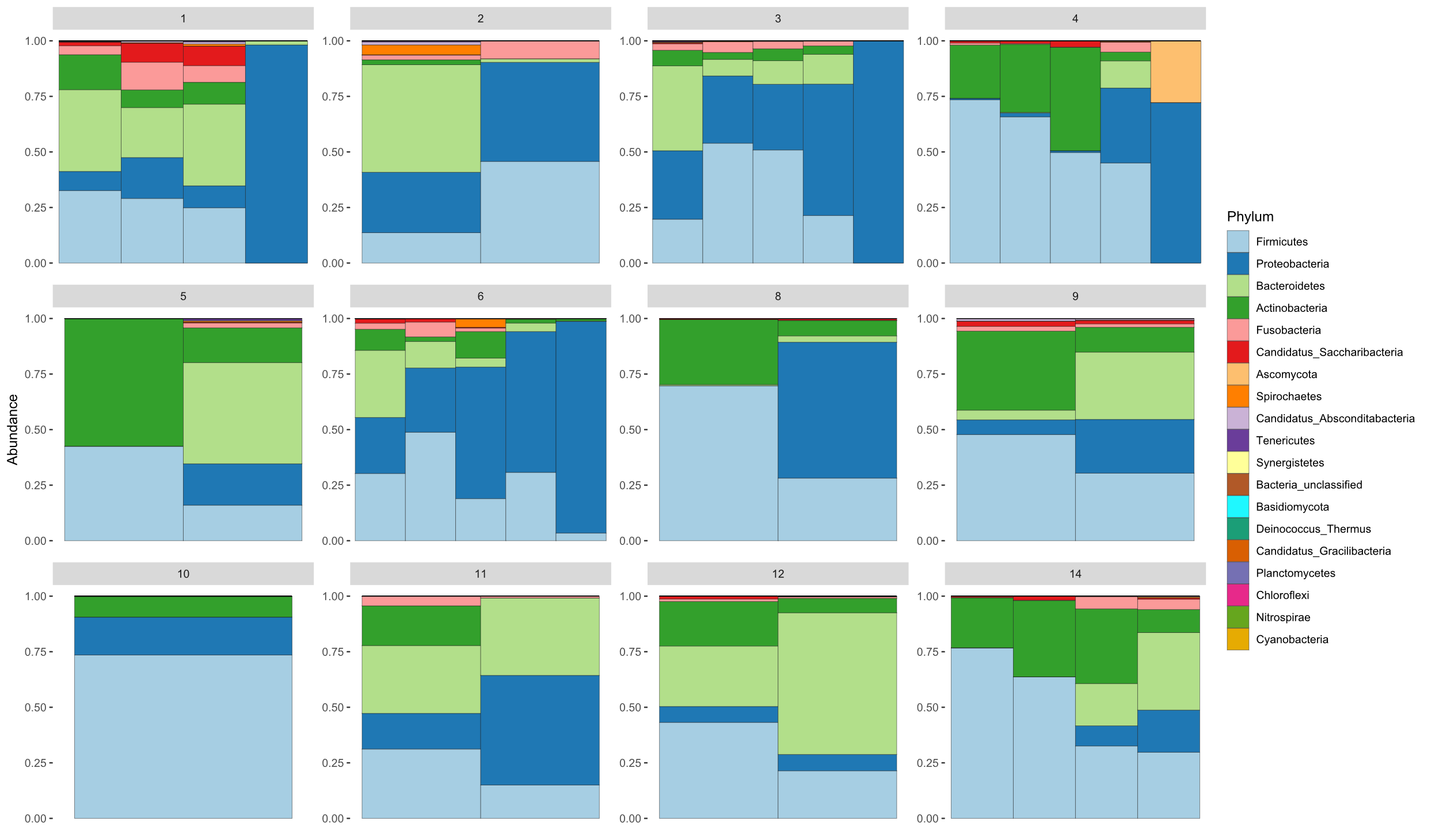

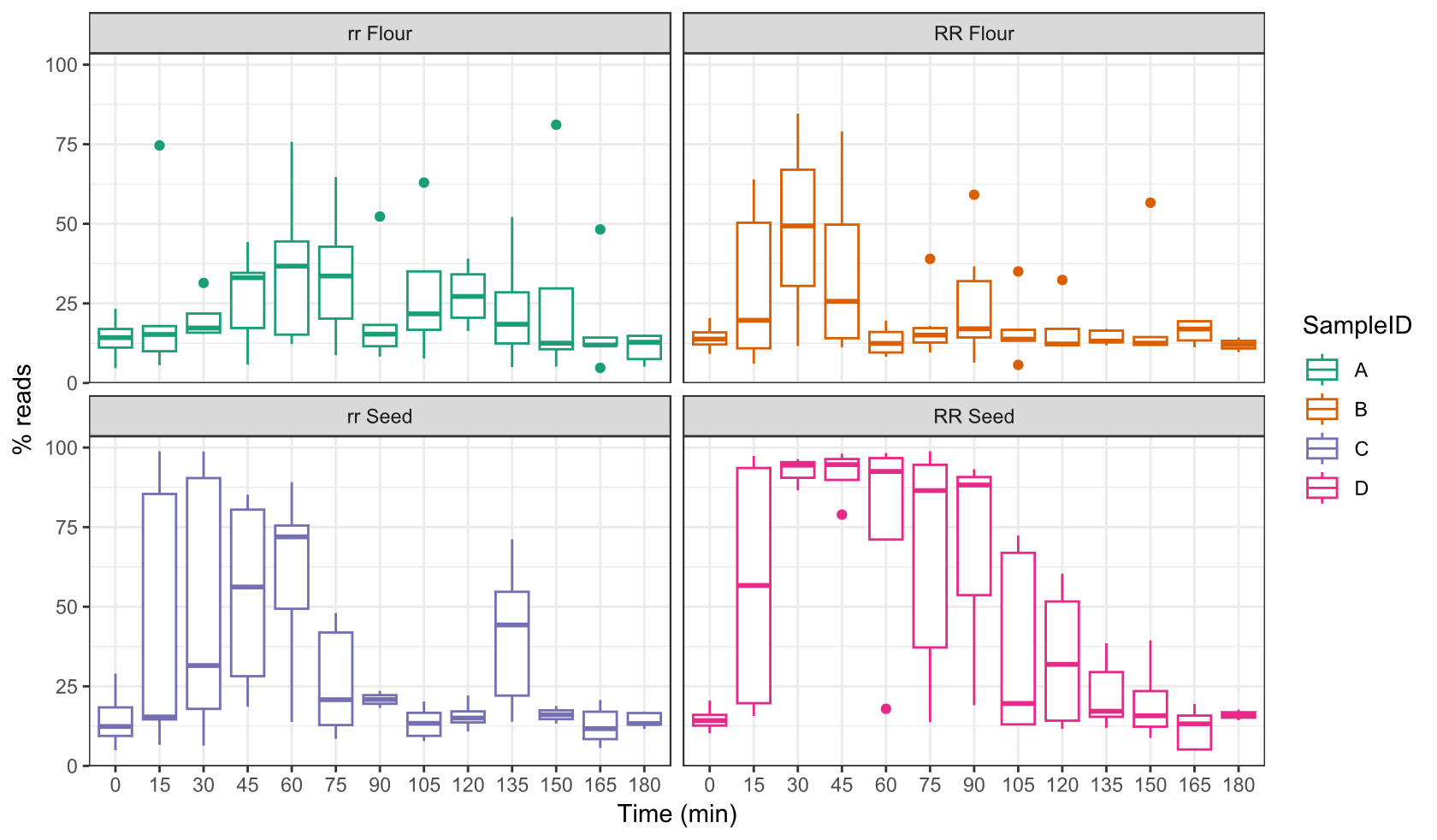

**Figure S1:** (A) Phylum level microbiome profiles of each participant for samples taken at baseline (B) Percentage of reads mapping to the Pisum Sativum genome at each time point sampled.

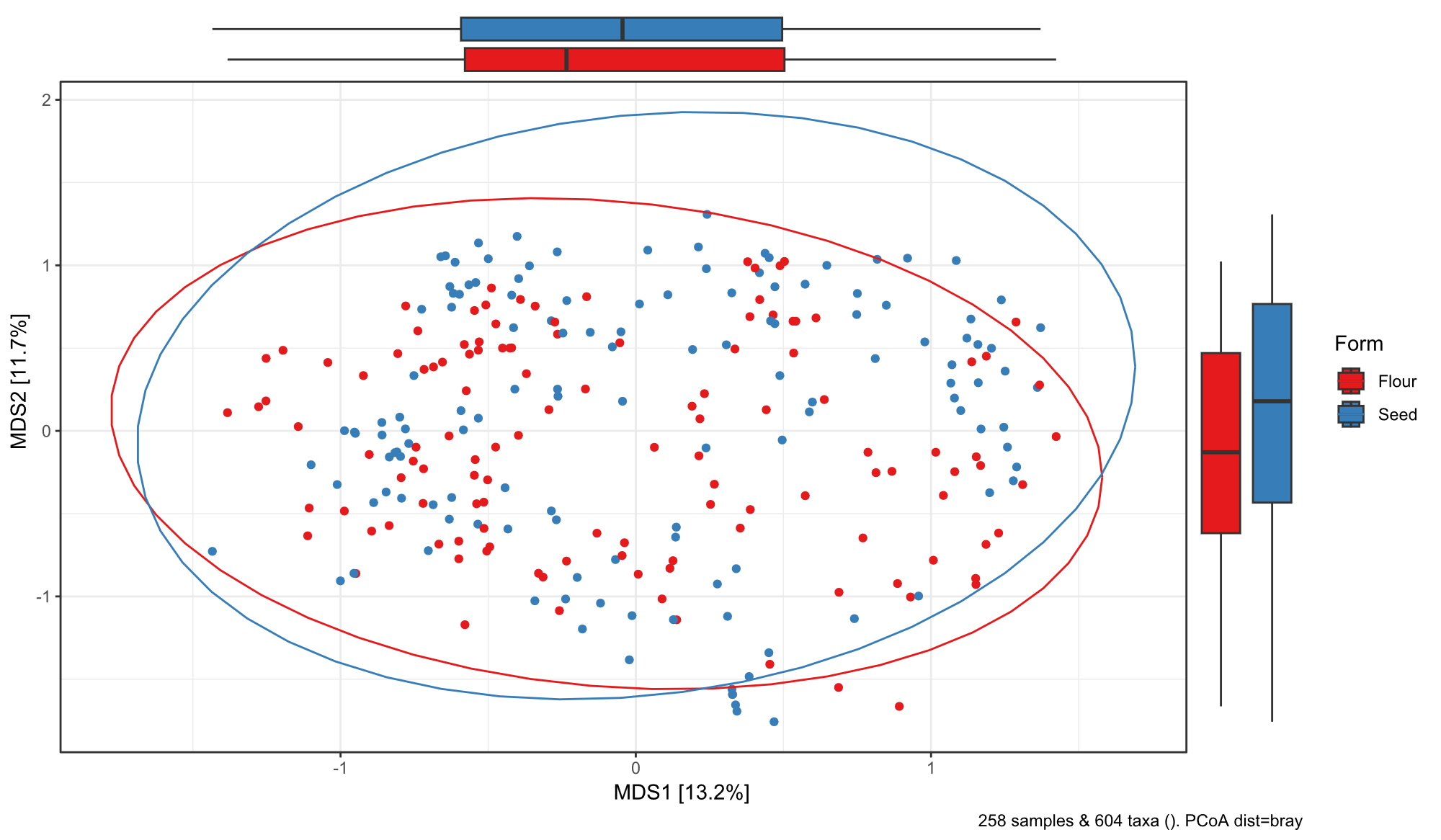

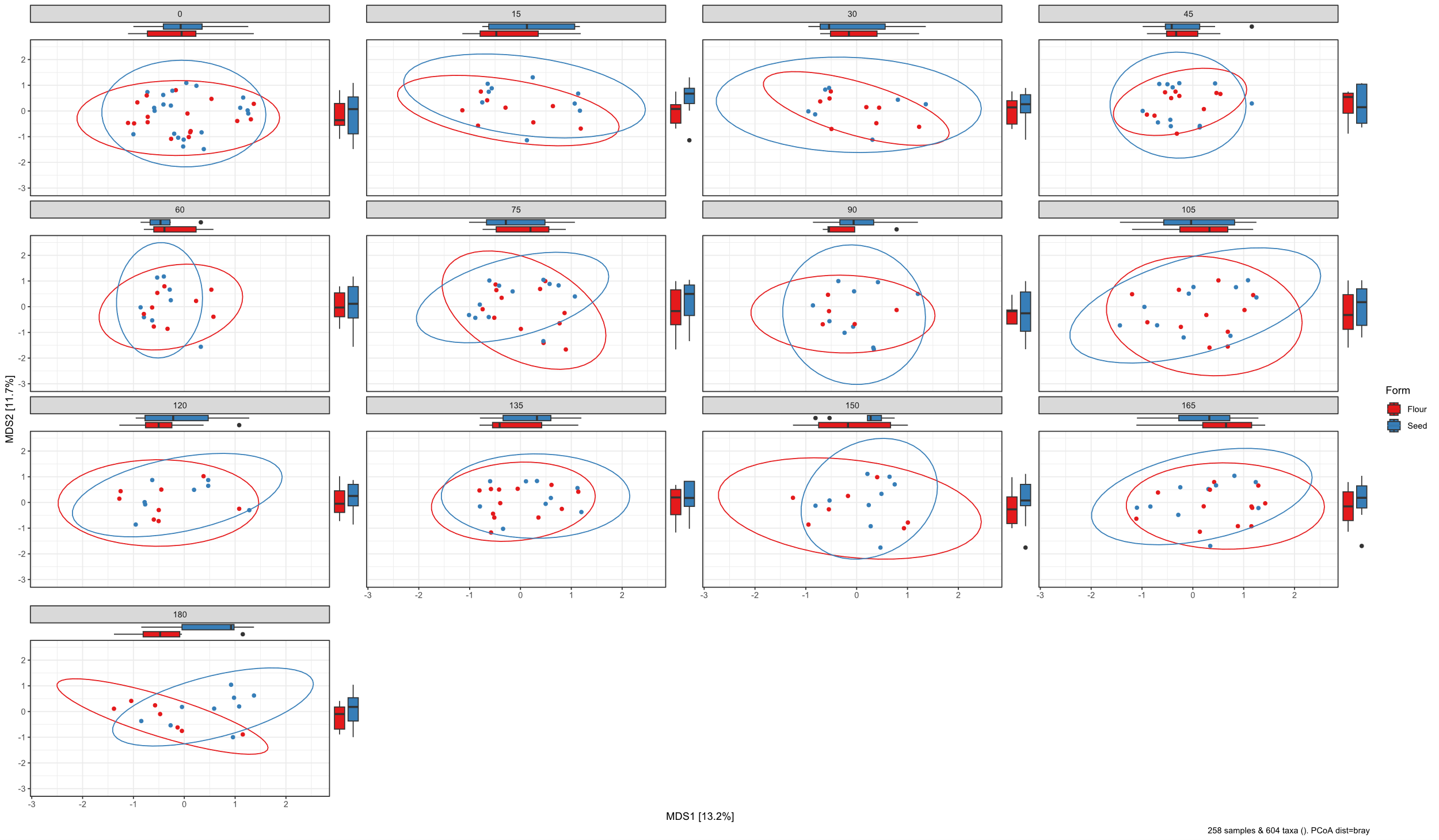

**Figure S2:** PCoA plot (Bray-Curtis dissimilarity) following consumption of the pea meals, coloured by the physical form of the pea meal (whole pea, vs. flour). **A.** showing all time points combined **B.** Showing community composition at each individual timepoint.

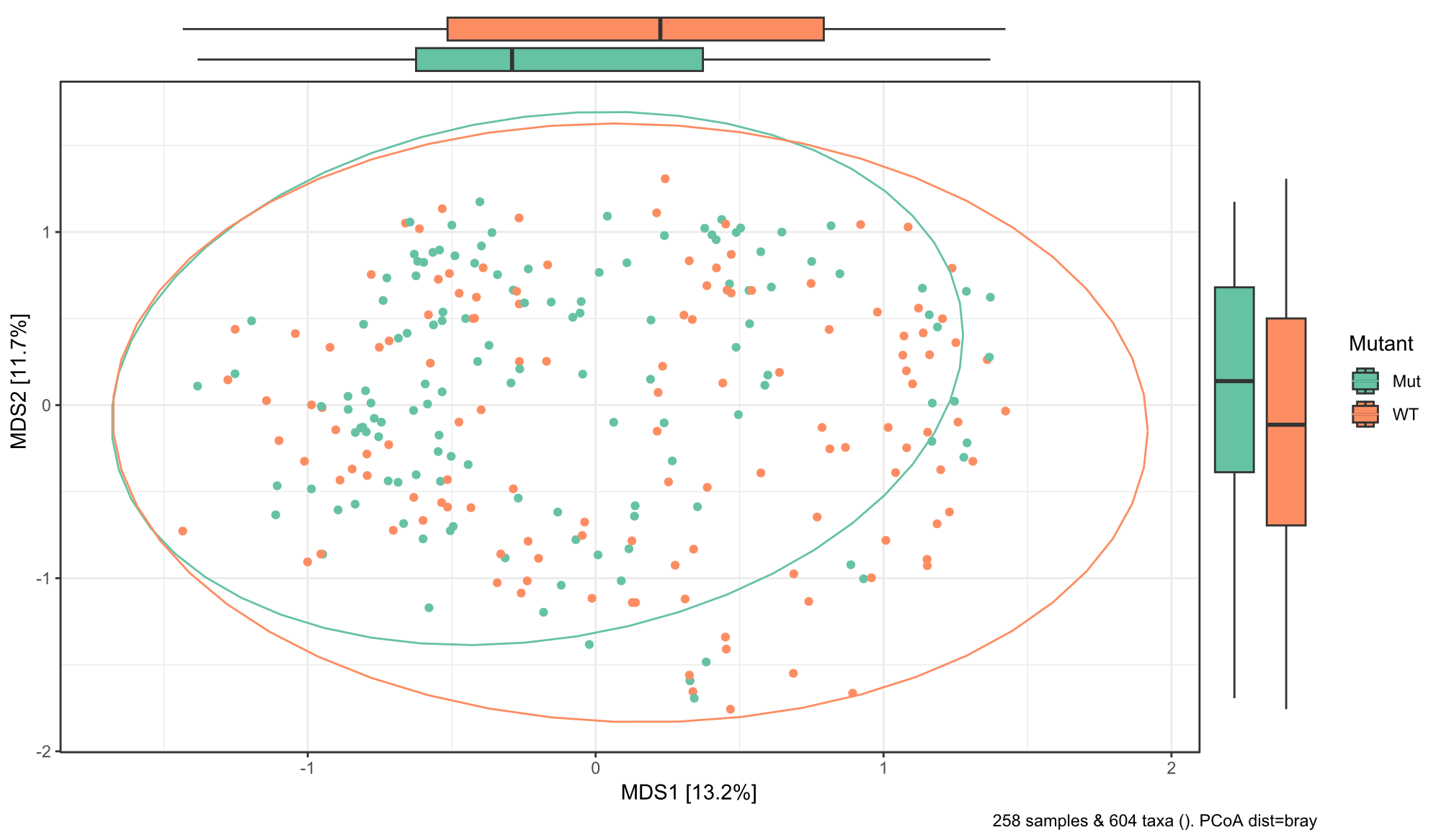

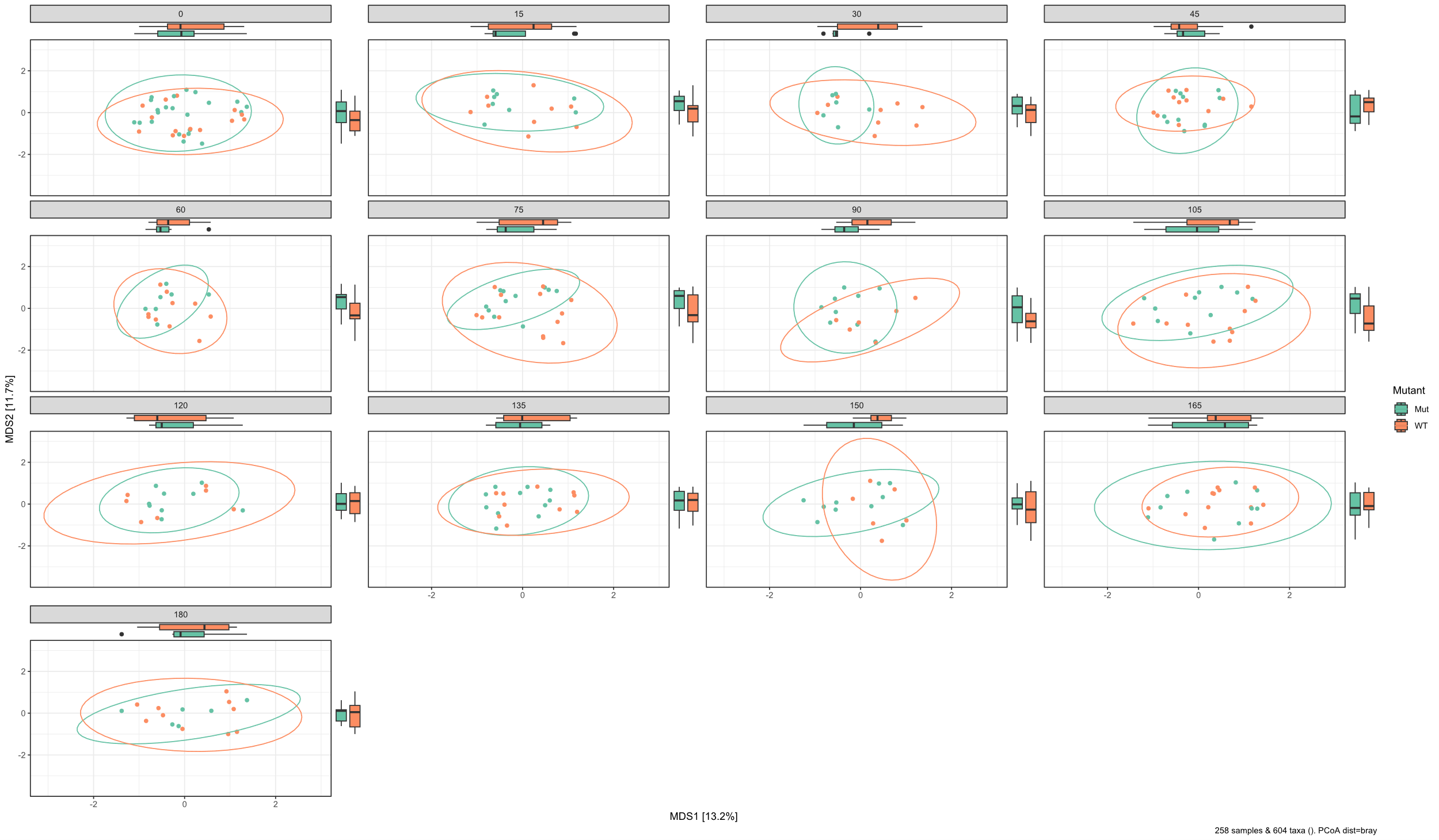

**Figure S3:** PCoA plot (Bray-Curtis dissimilarity) following consumption of the pea meals, coloured by the genotype of the flour **A.** showing all time points combined **B.** Showing community composition at each individual timepoint.

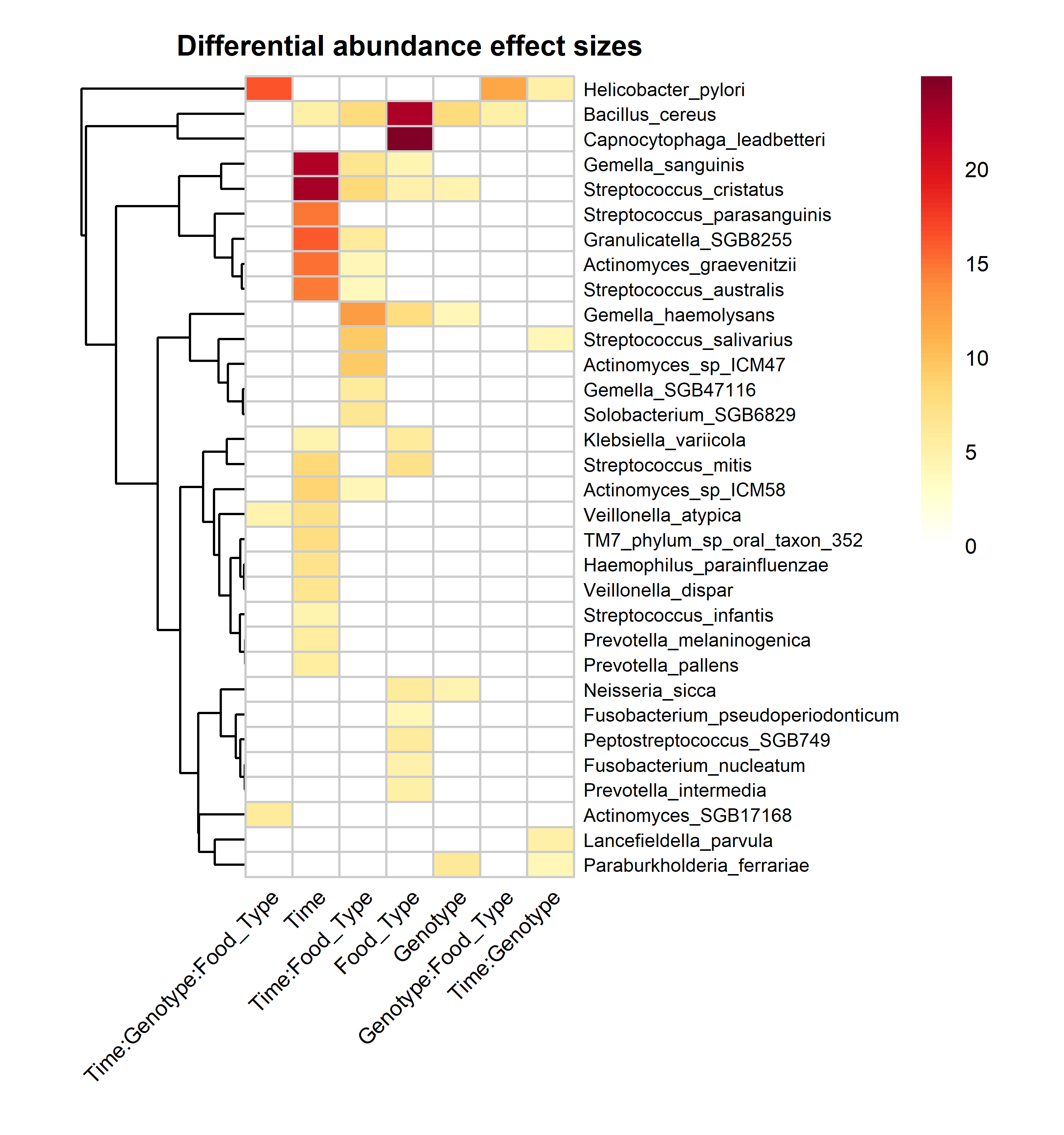

**Figure S4: Effect size heatmap for differential abundance.** Rows represent individual bacterial taxa that showed significant differential abundance (P < 0.05) for at least one experimental factor. Columns represent the experimental effects tested: main effects (Time, Genotype, Food_Type) and all two-way and three-way interactions. Cell color intensity indicates F-value from linear mixed-effects model ANOVA, with darker red indicating stronger effects. Rows are hierarchically clustered (Euclidean distance). White cells indicate non-significant effects (P ≥ 0.05) for that taxon-effect combination.

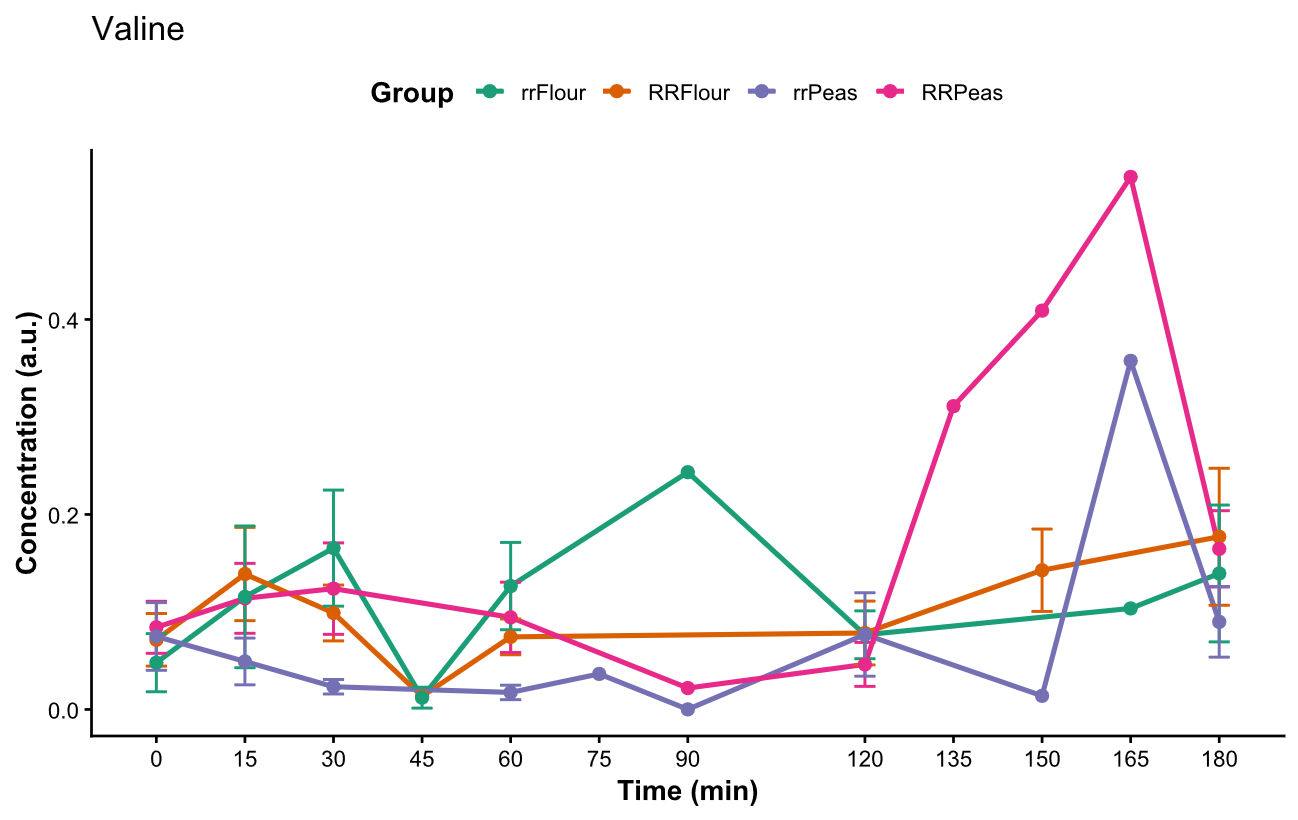

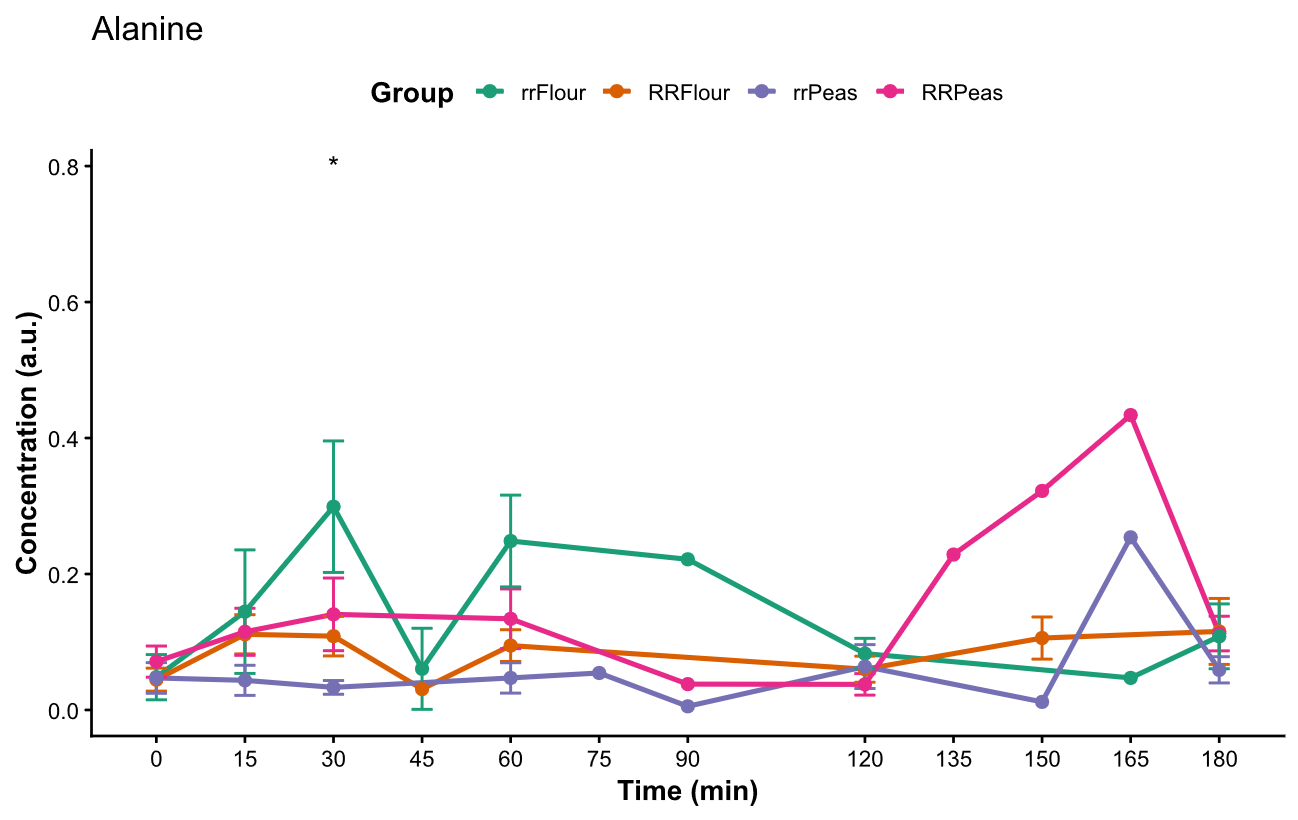

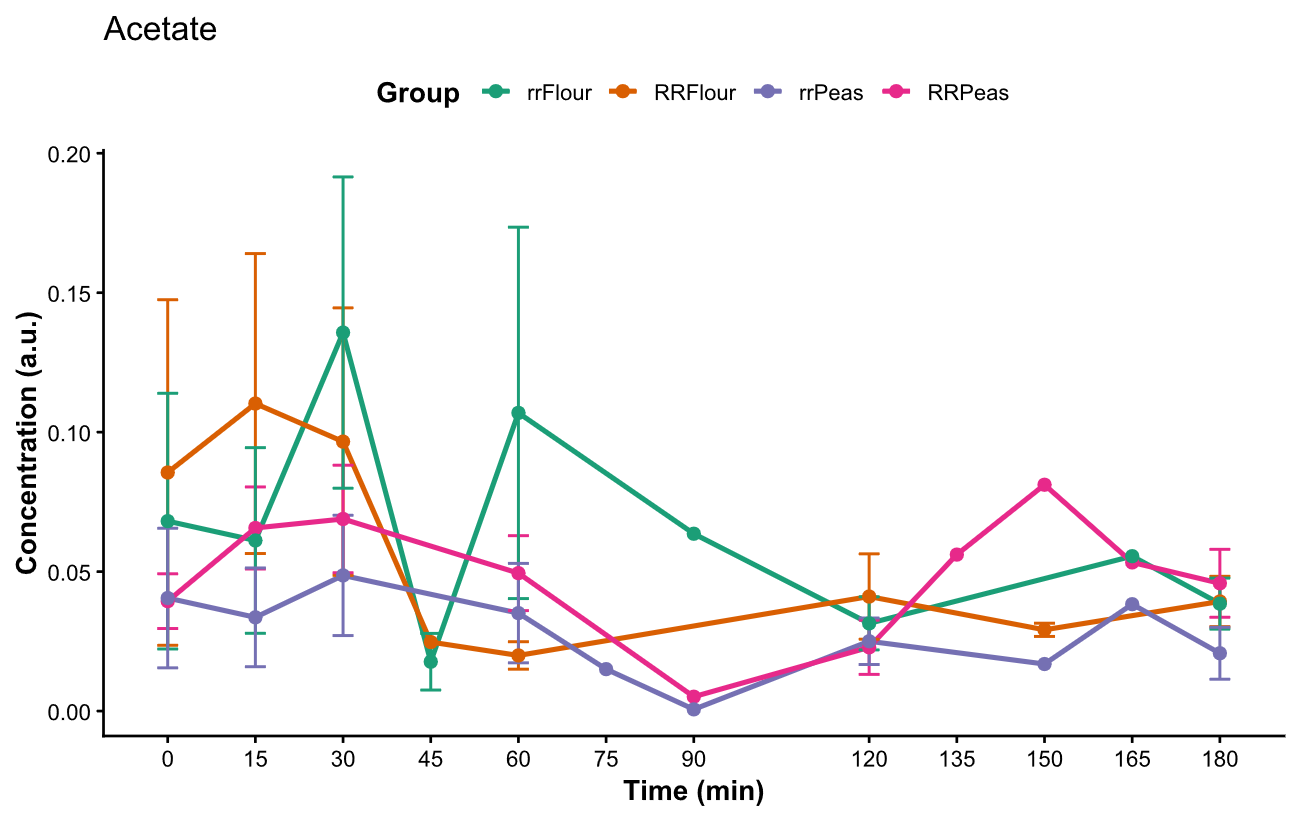

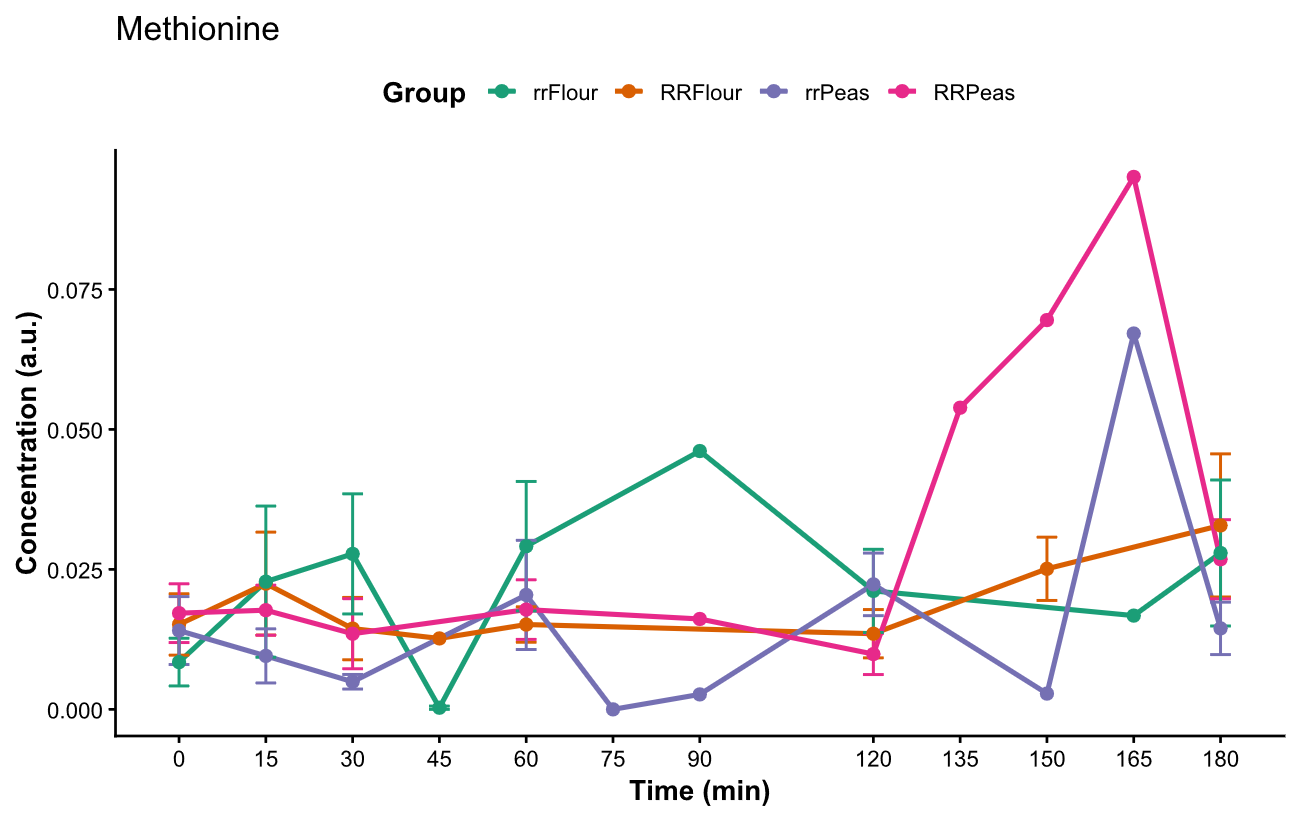

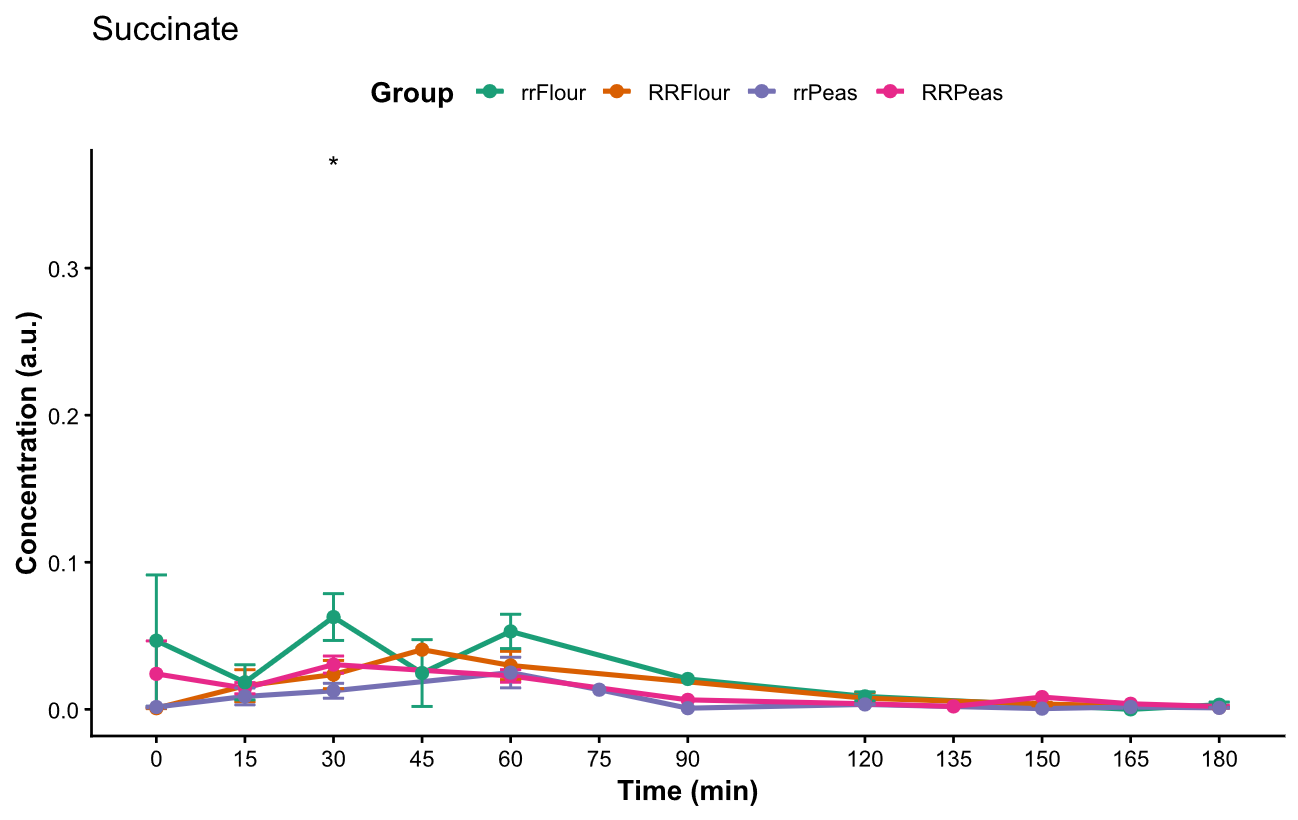

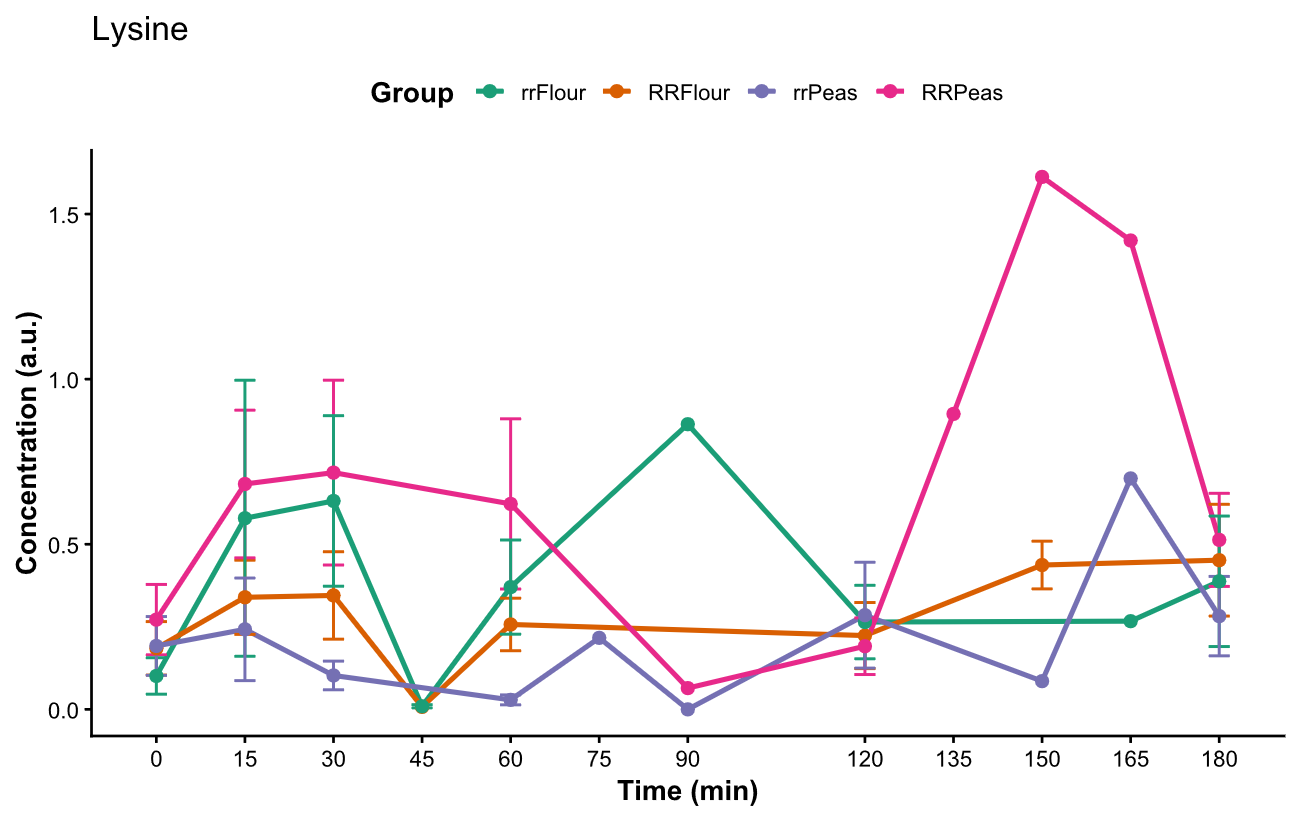

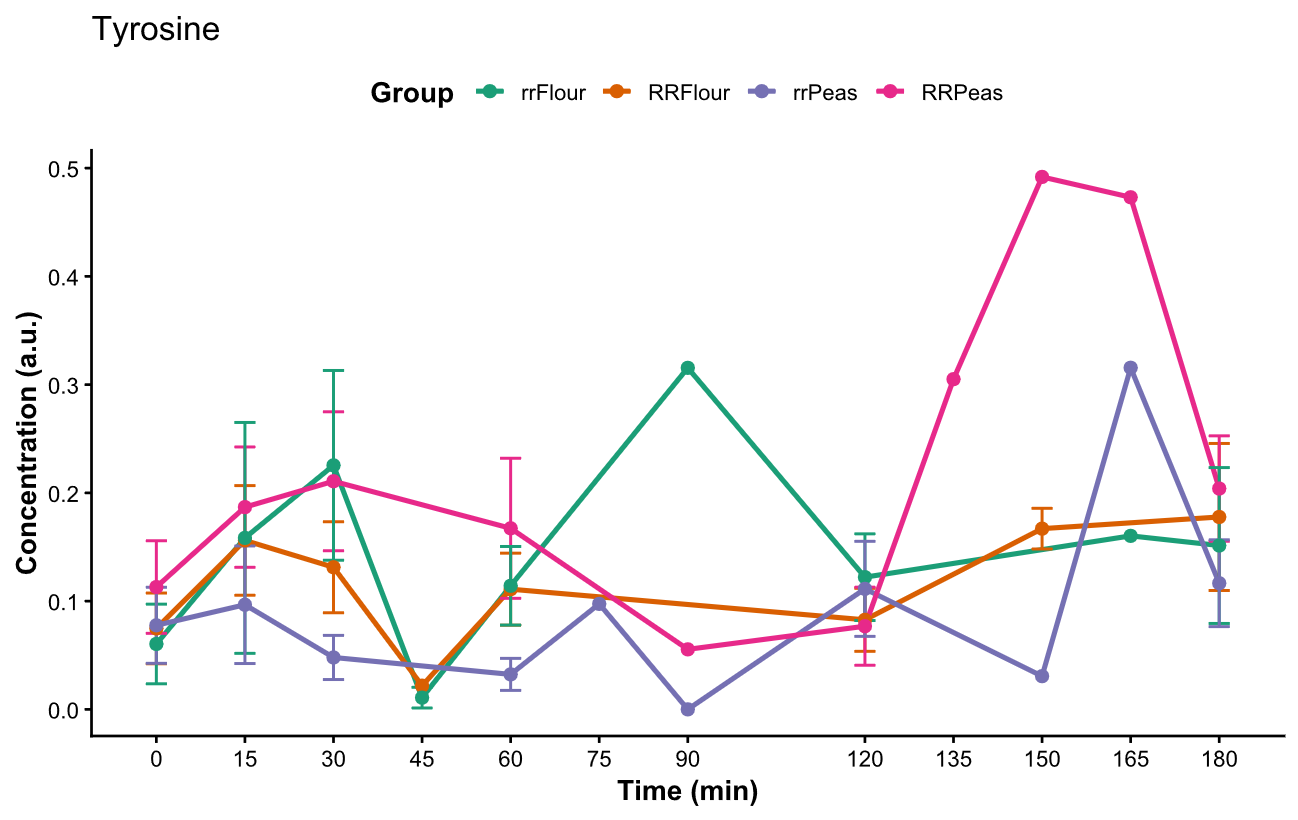

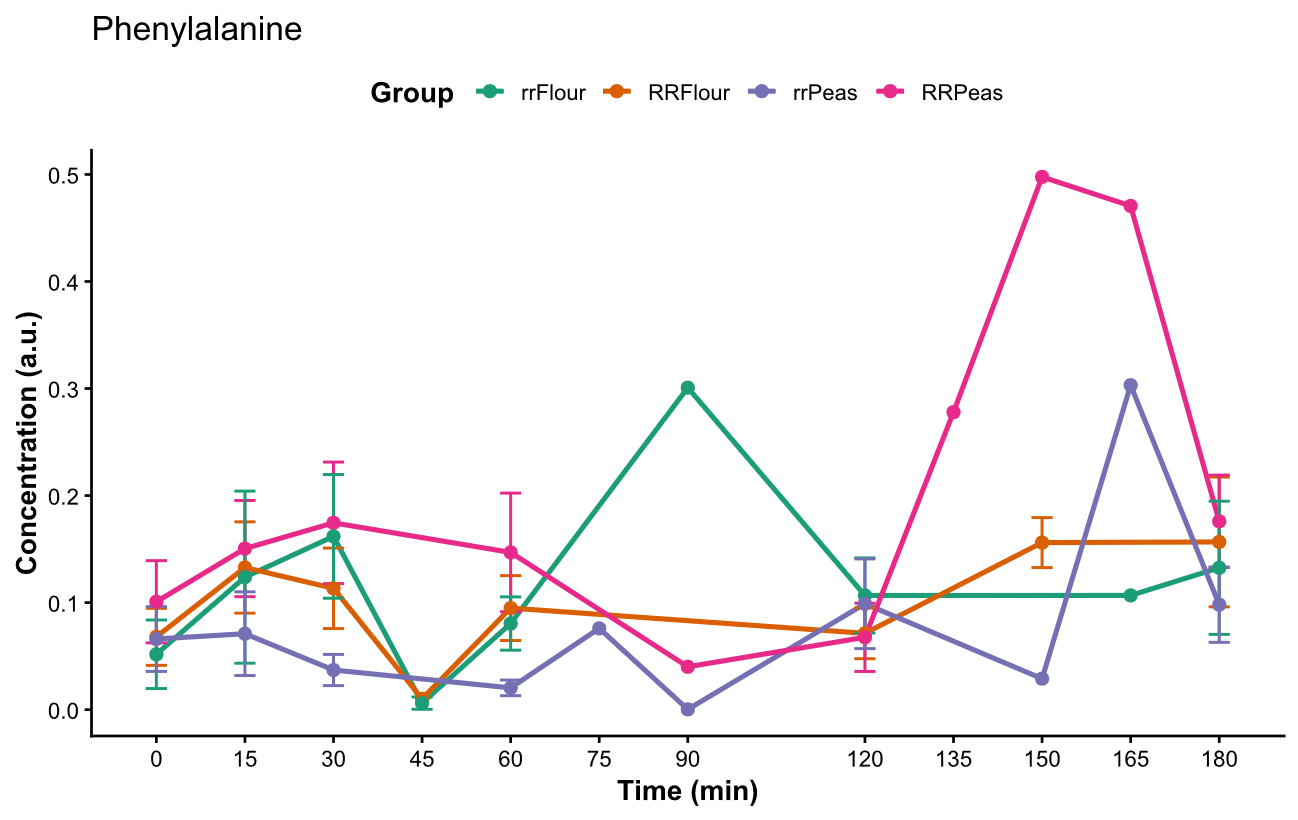

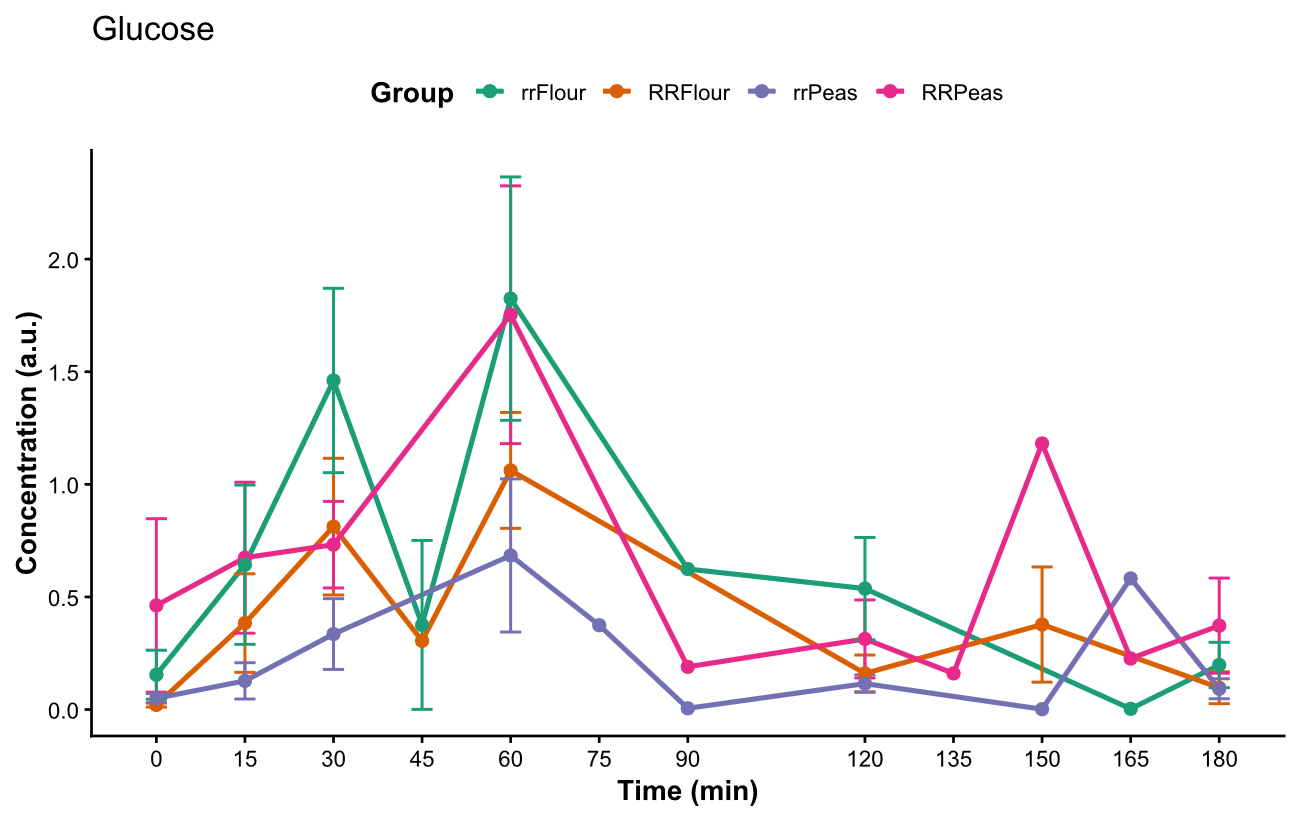

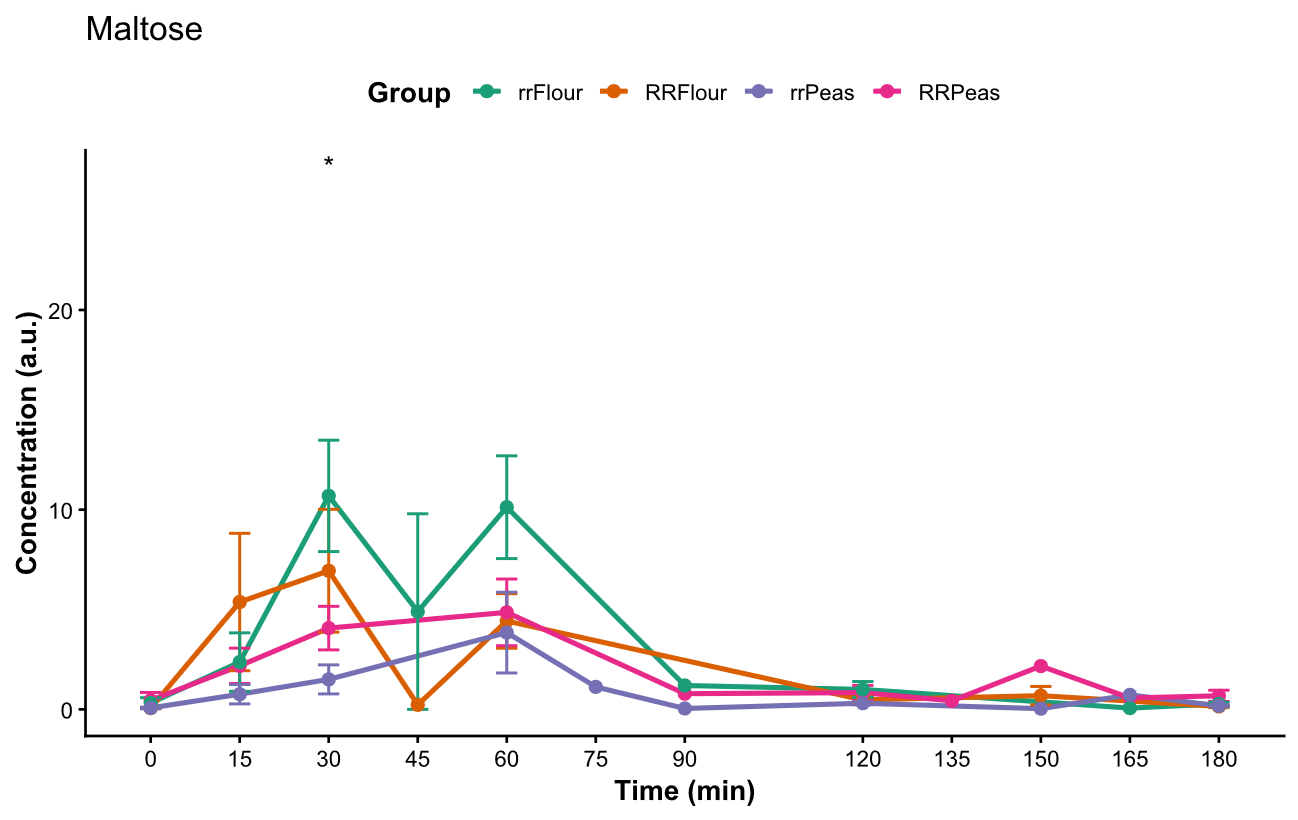

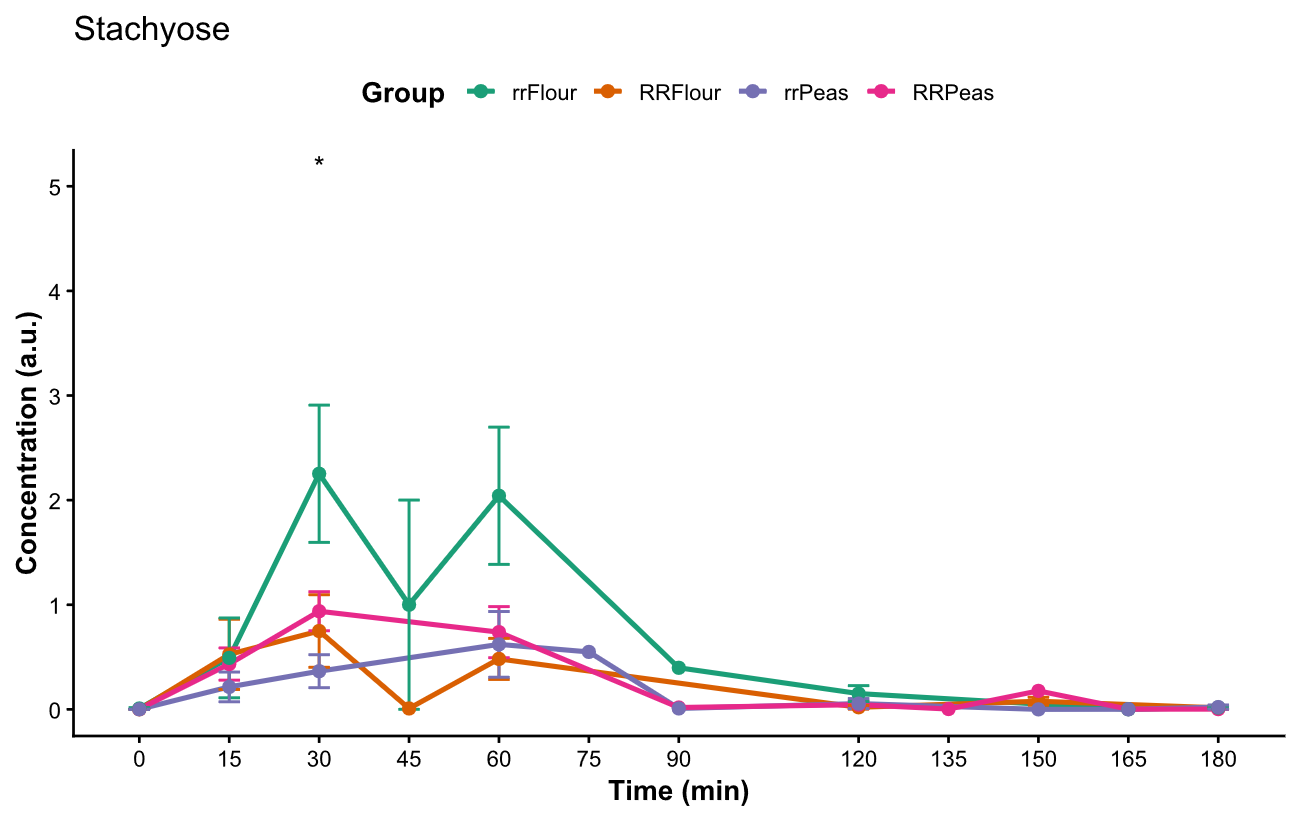

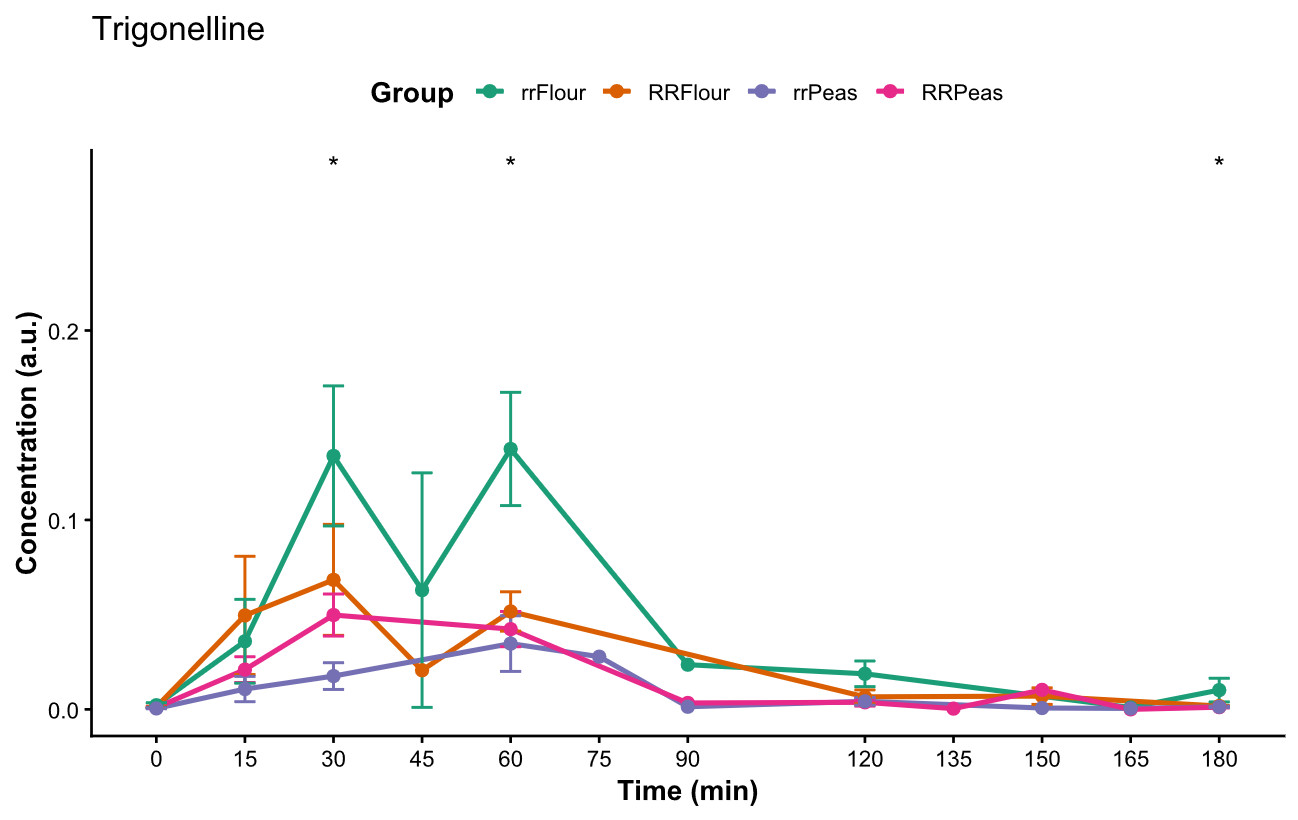

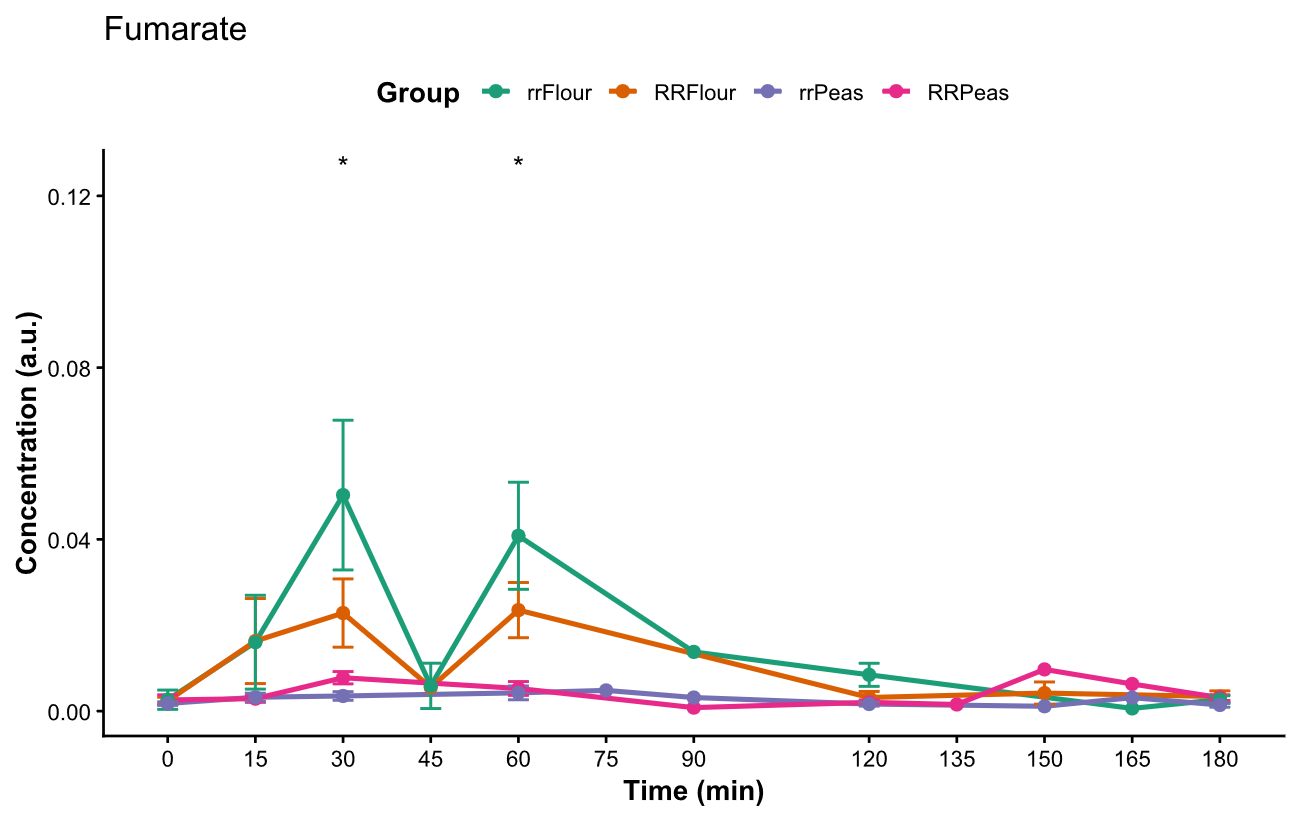

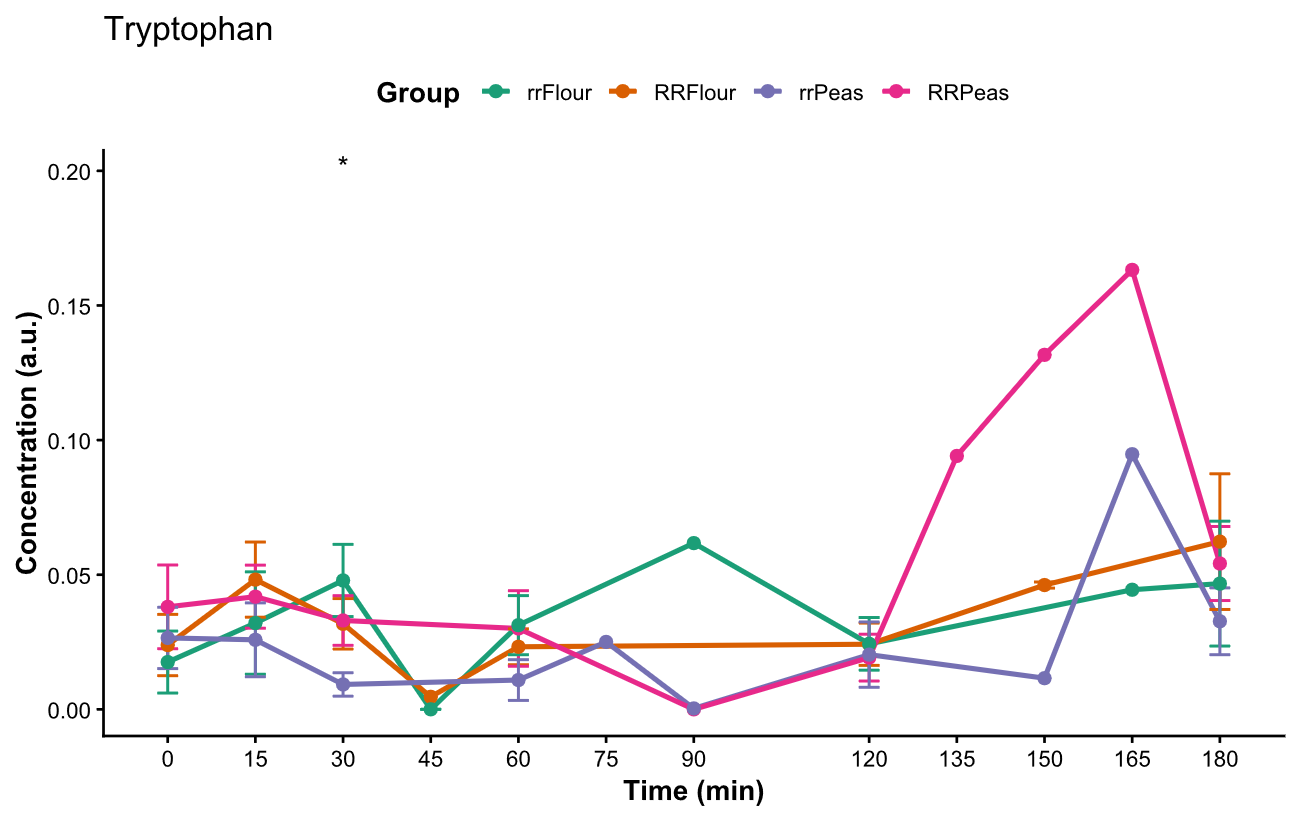

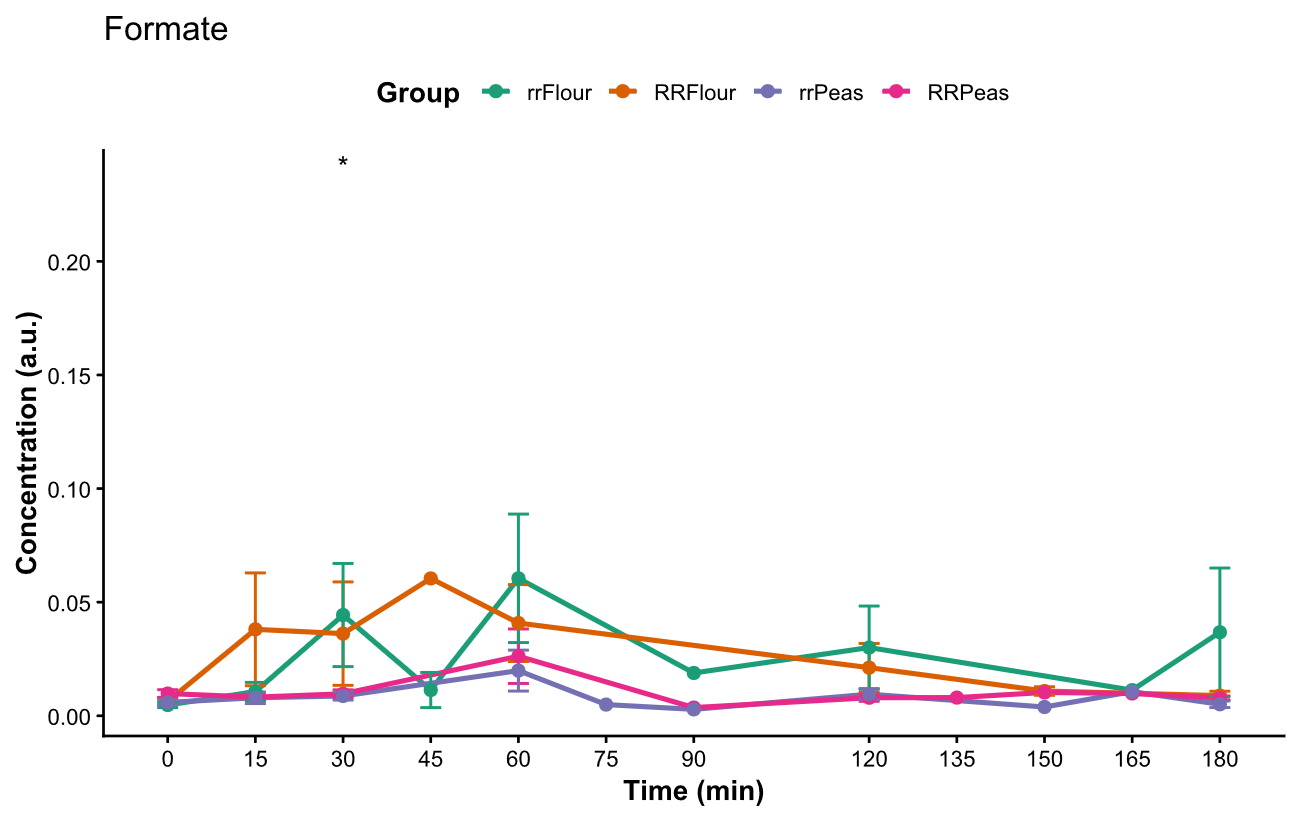

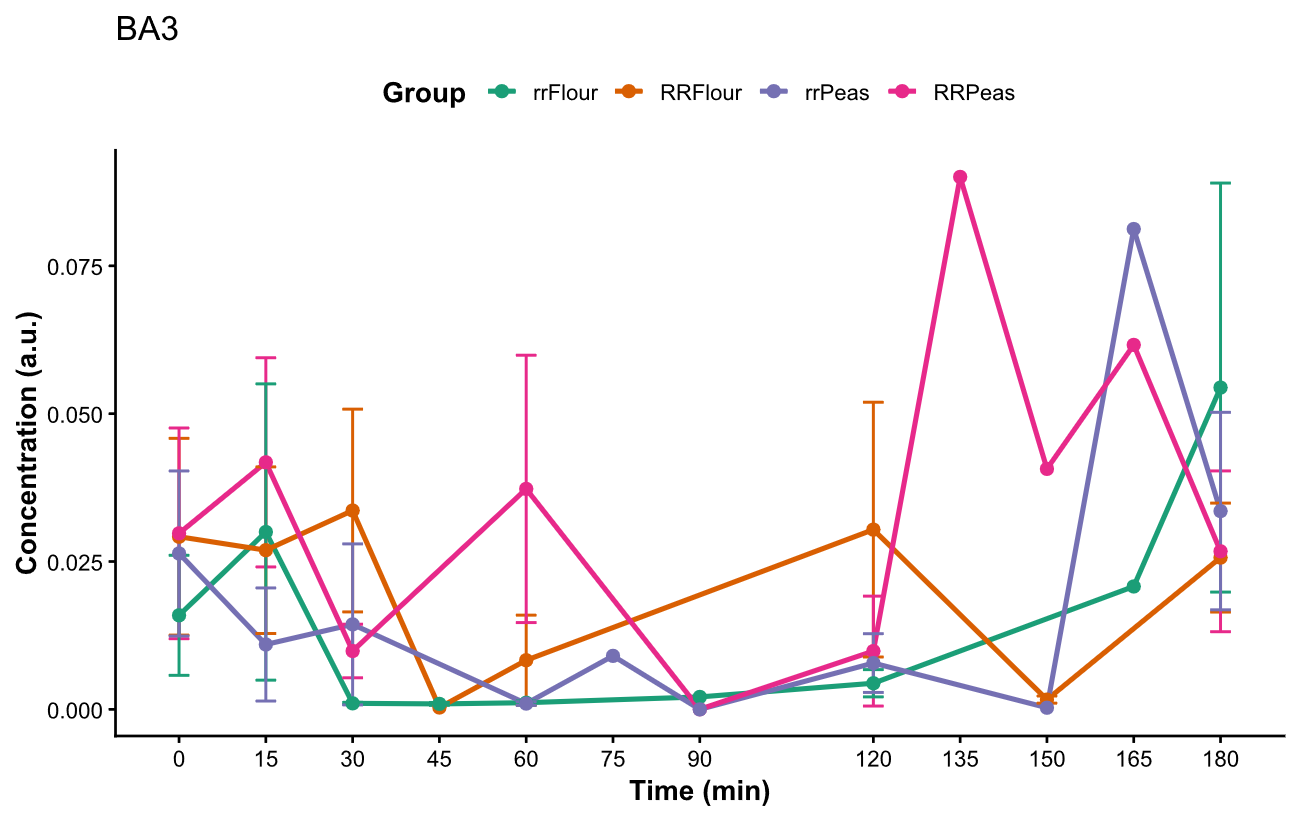

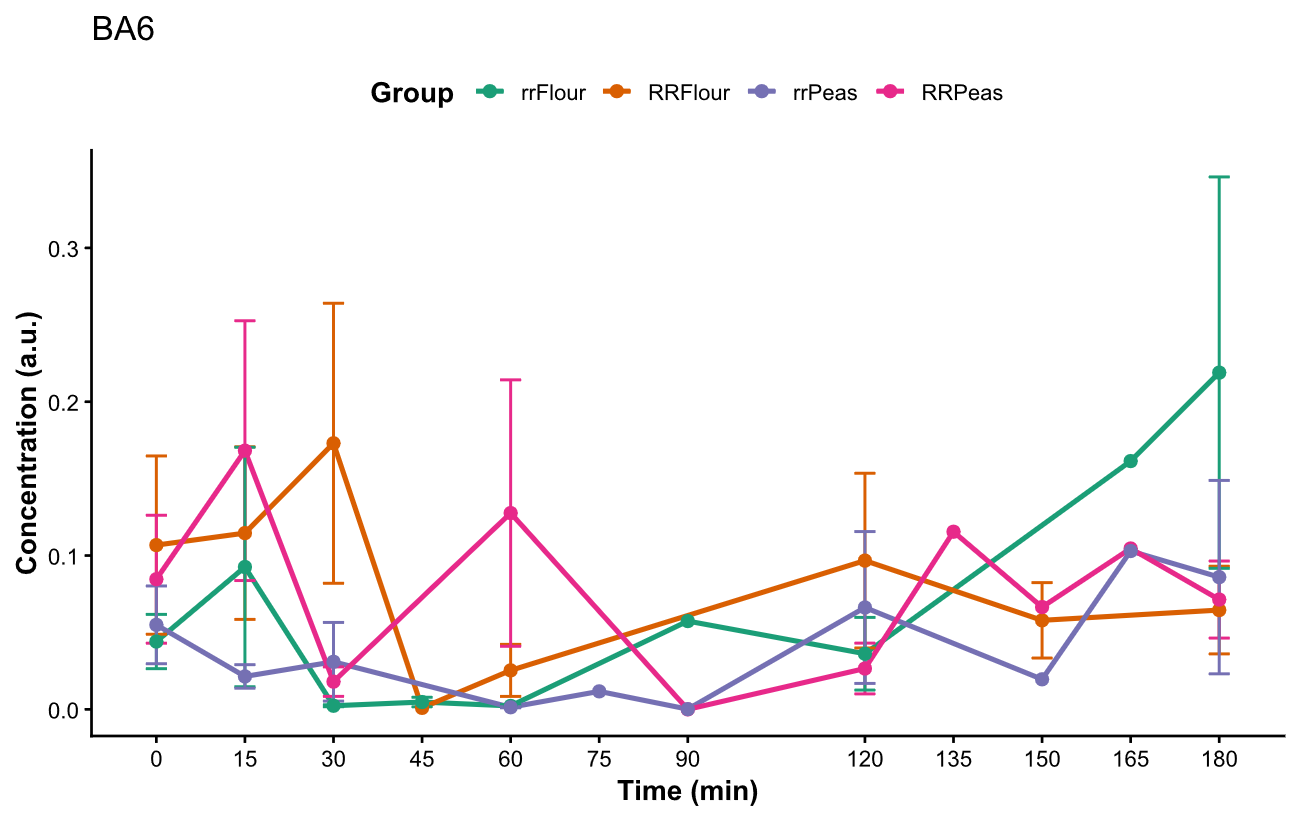

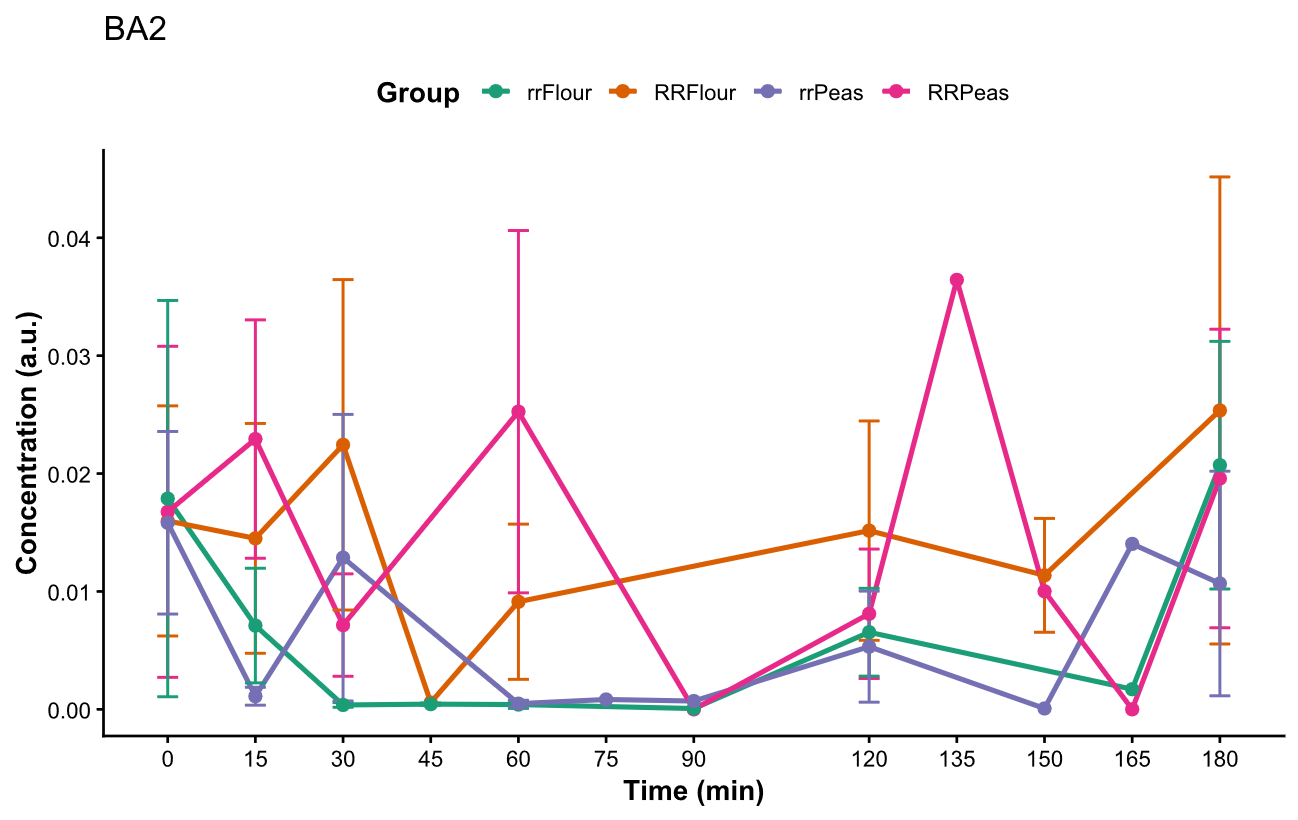

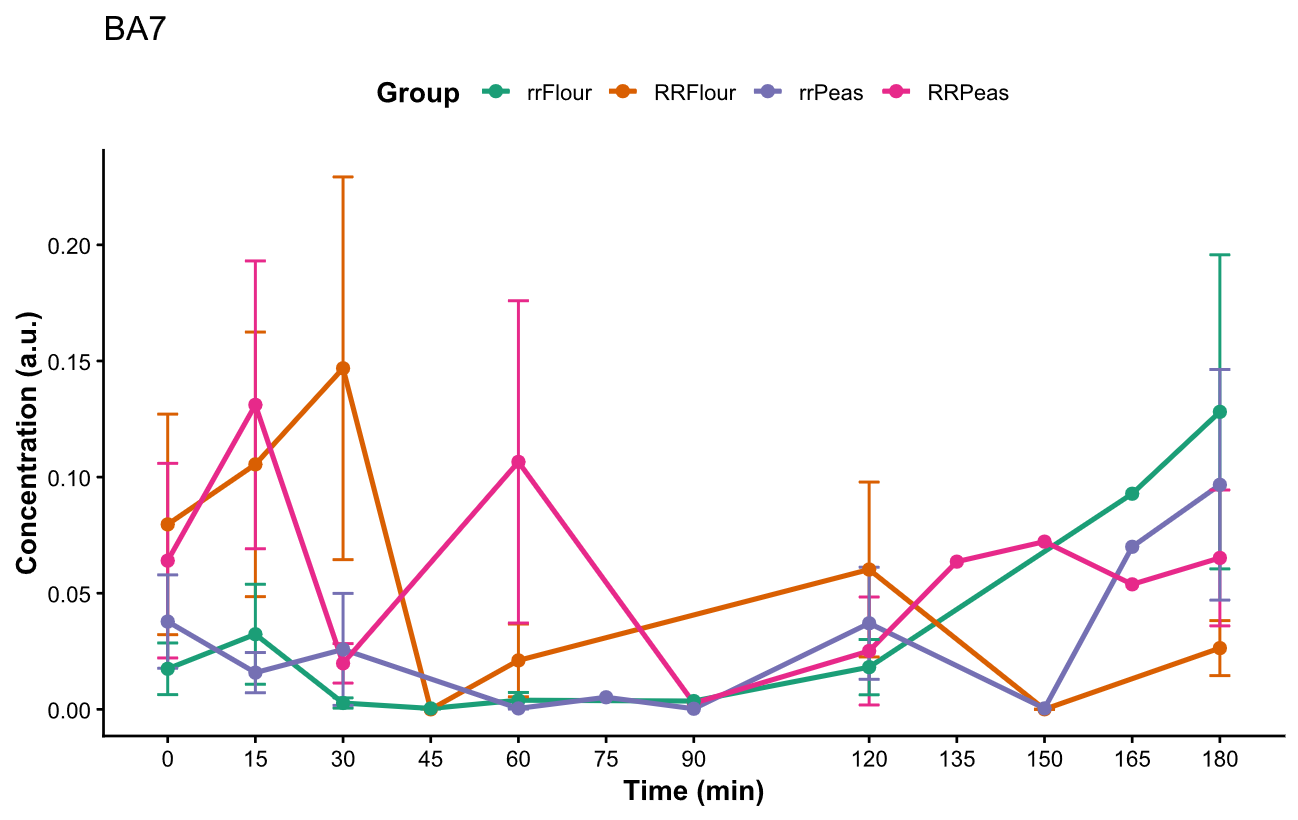

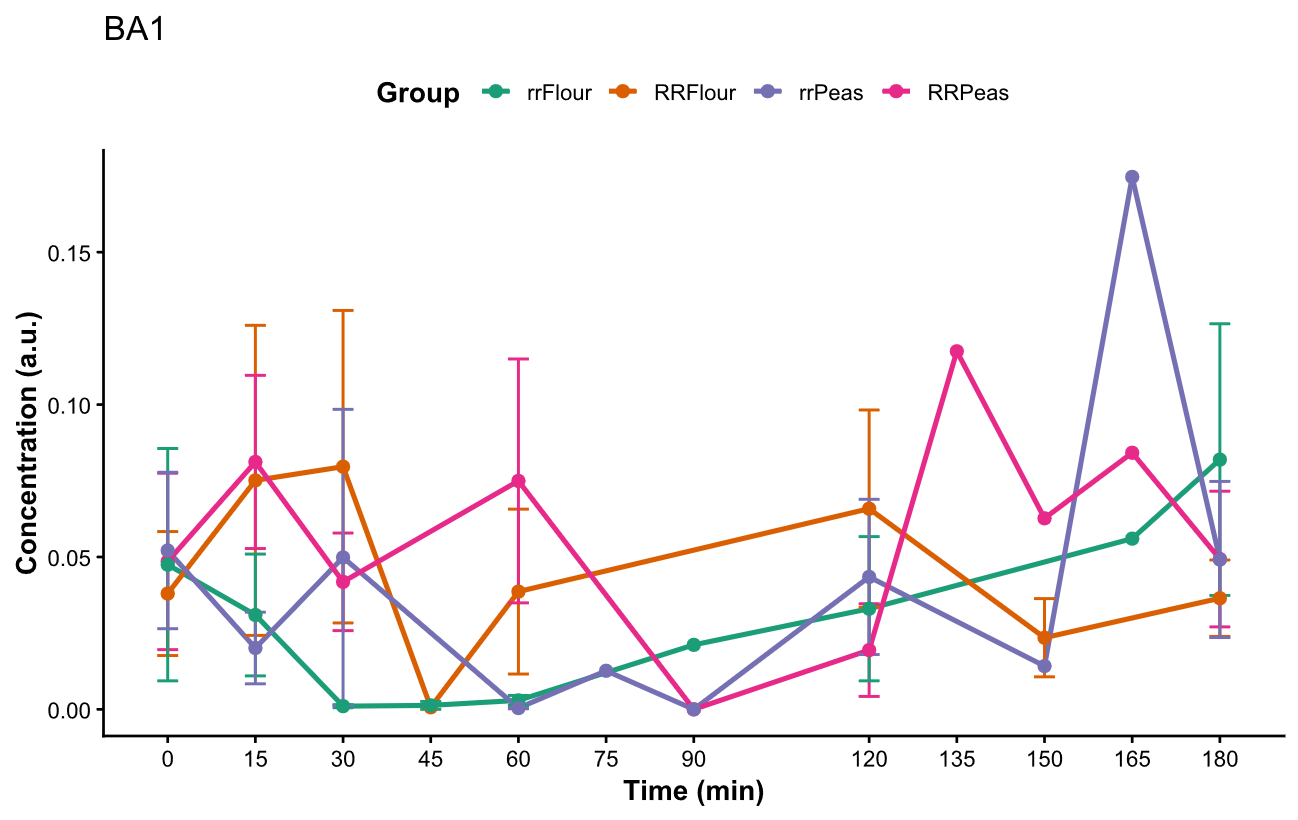

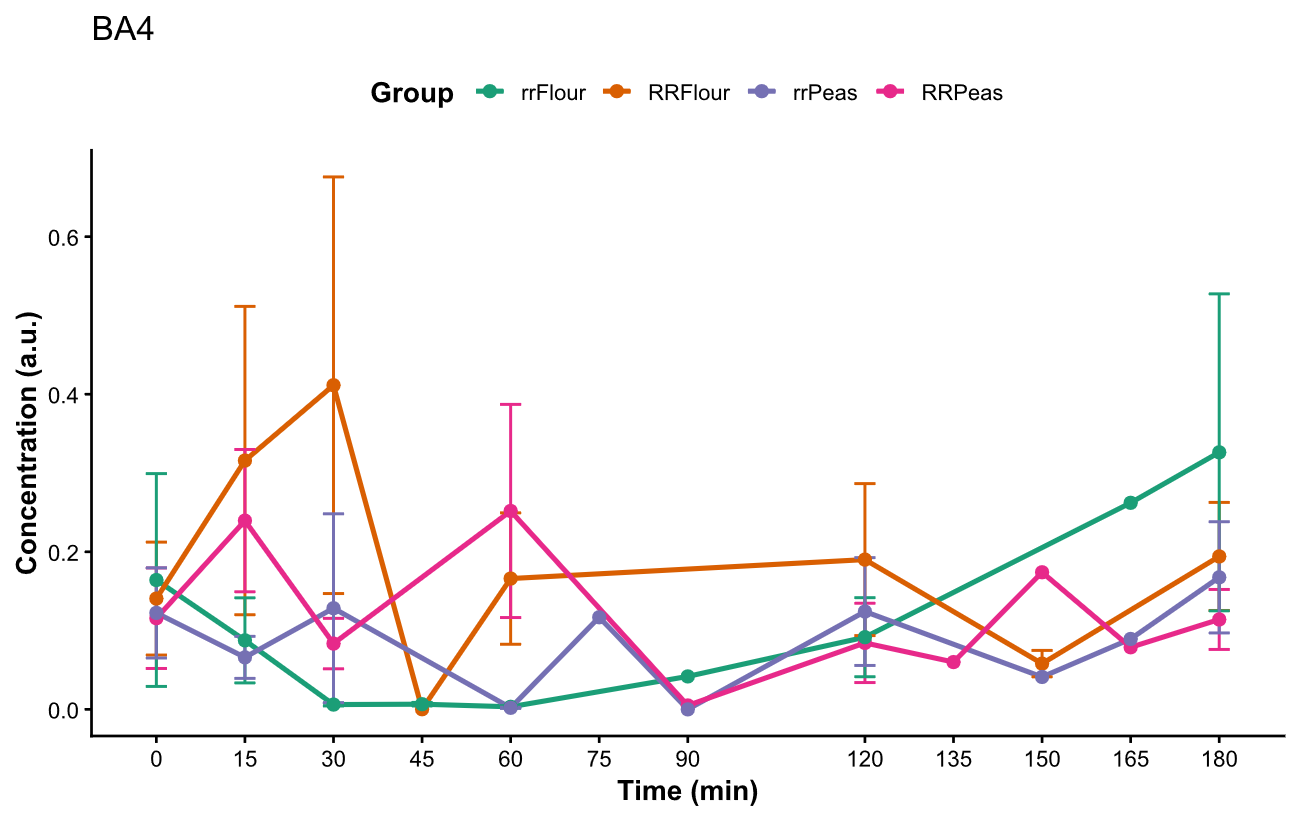

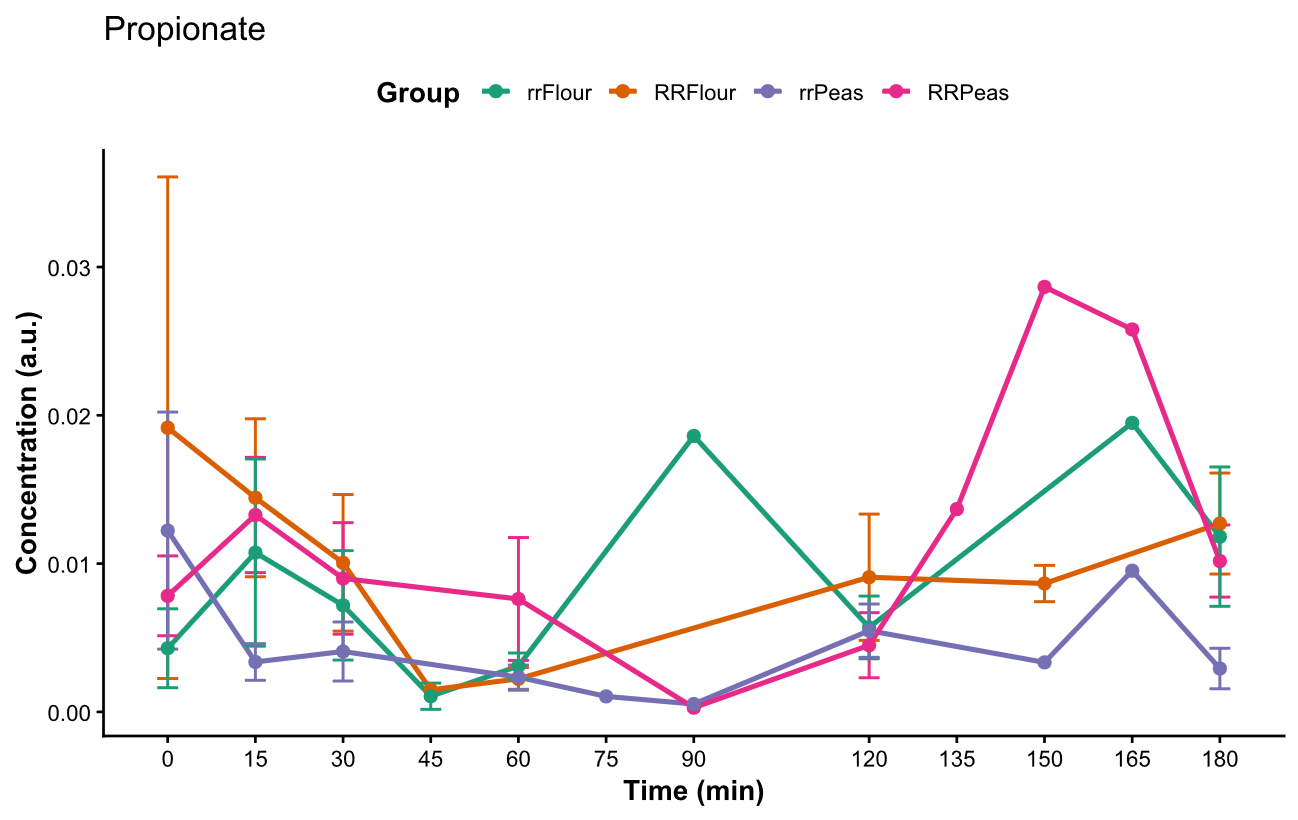

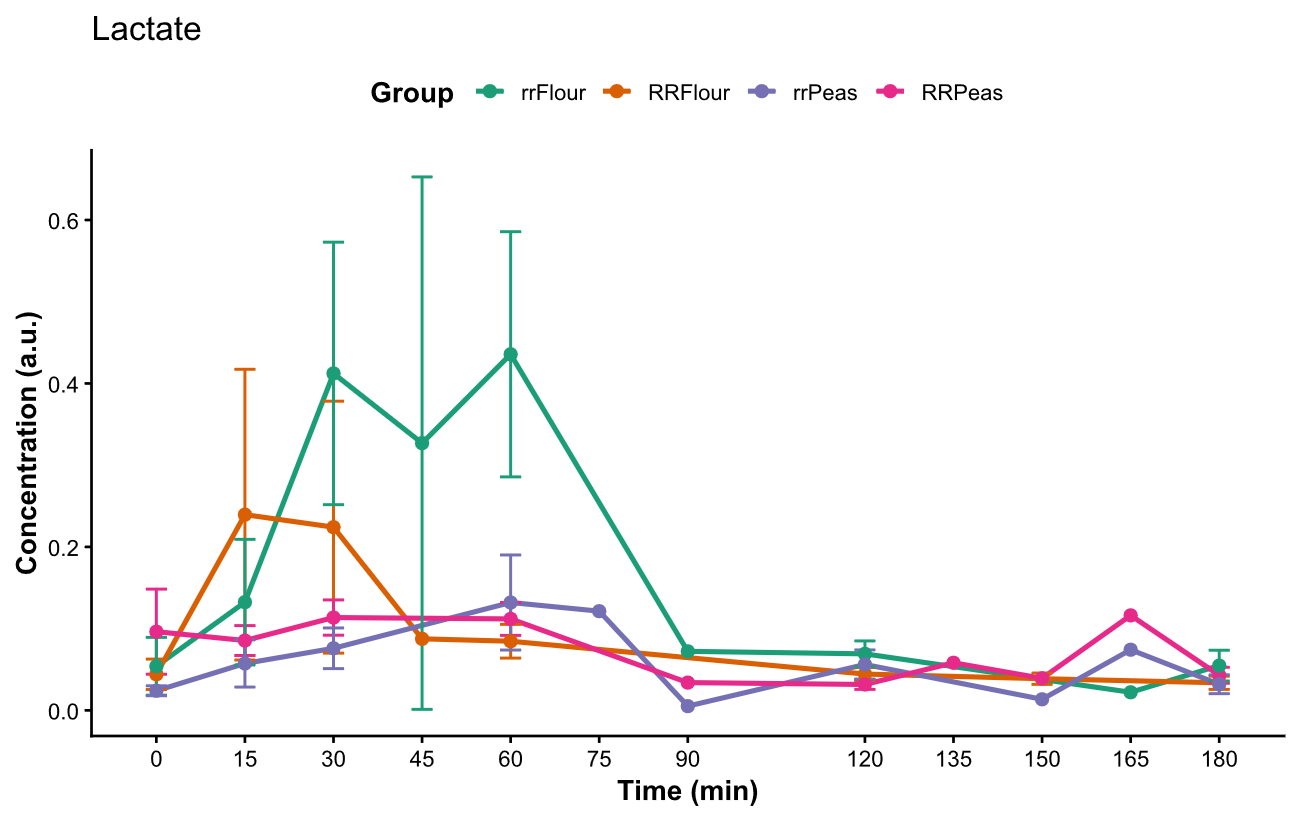

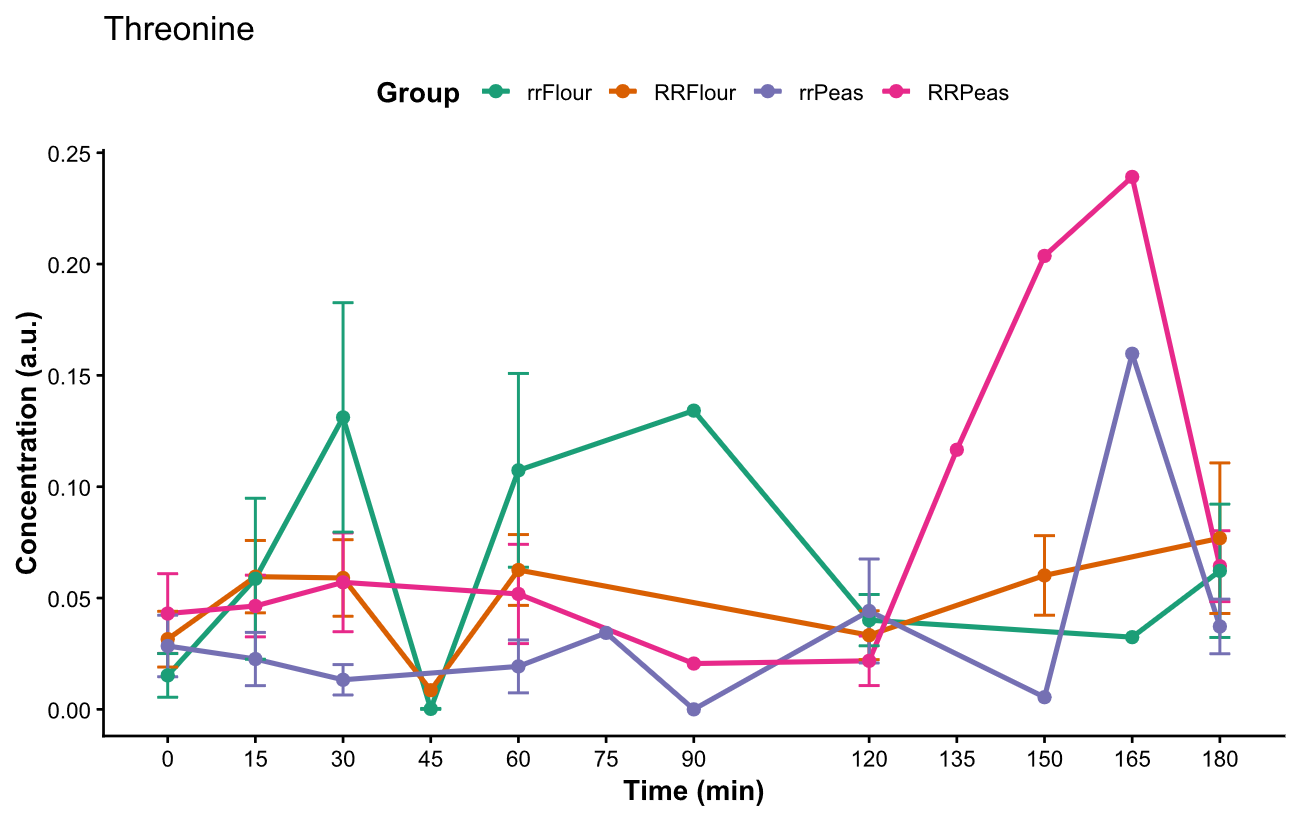

**Figure S5: Metabolite profiles over time measured by 1H NMR spectroscopy in duodenal samples.** Each of the profiles are coloured by meal type. Significant differences (Kruskal-Wallis) between meal types are indicated with *.

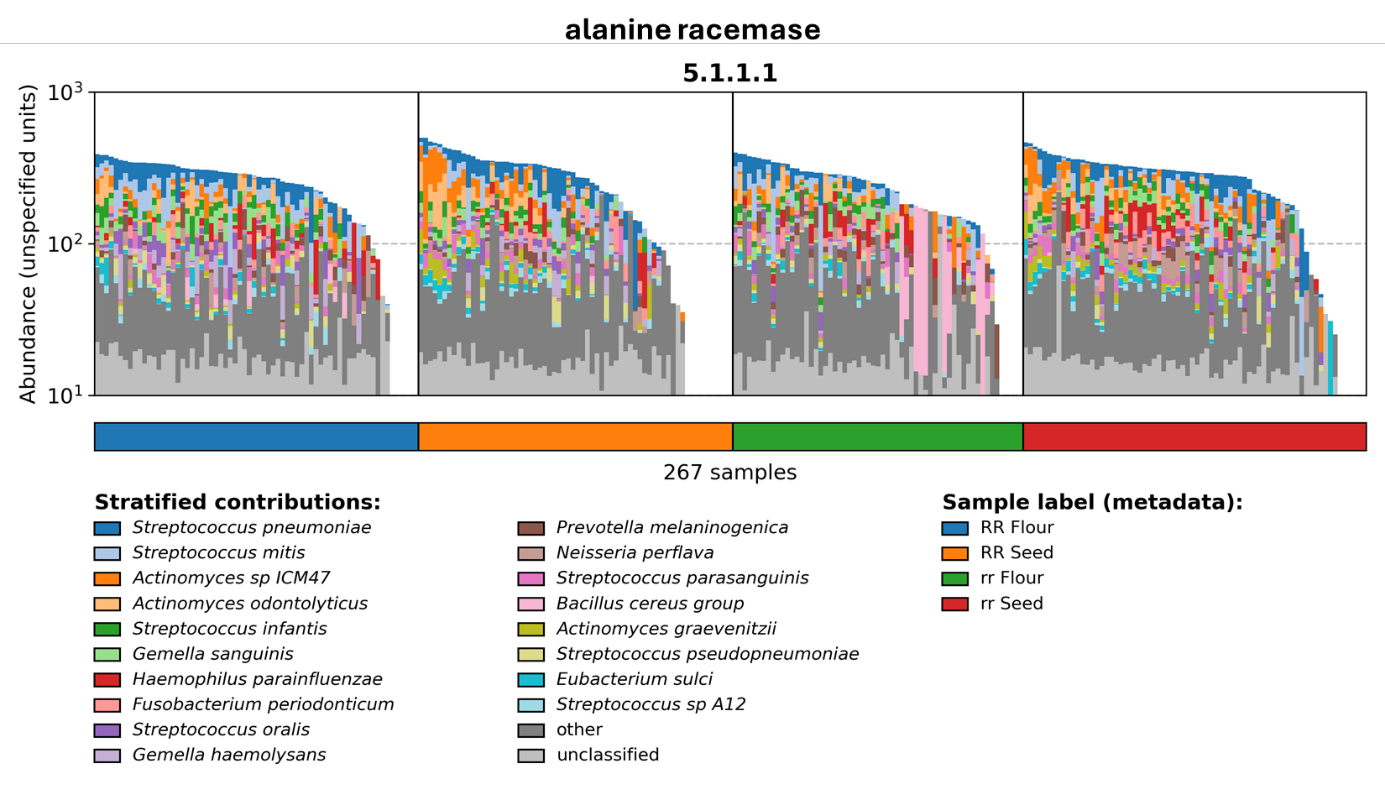

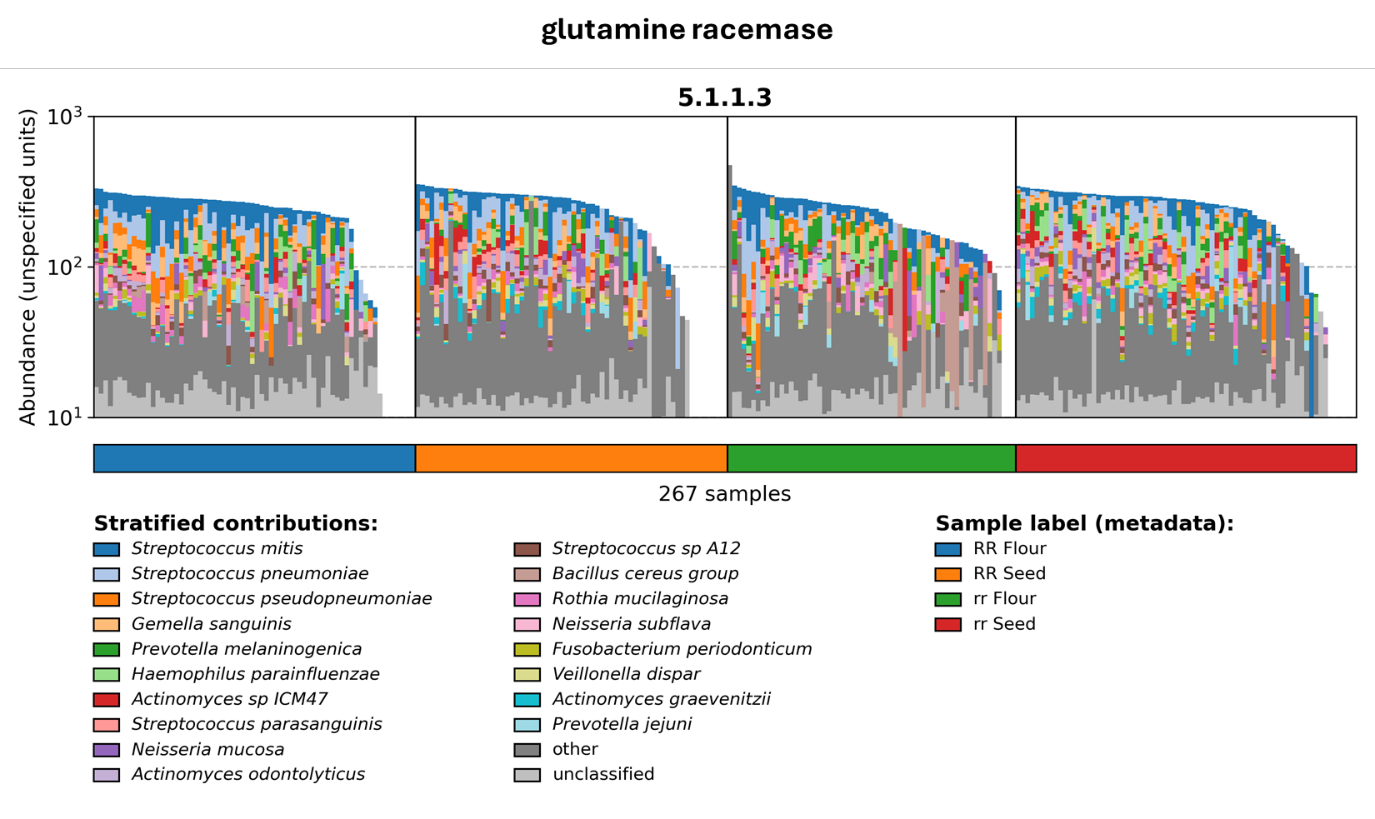

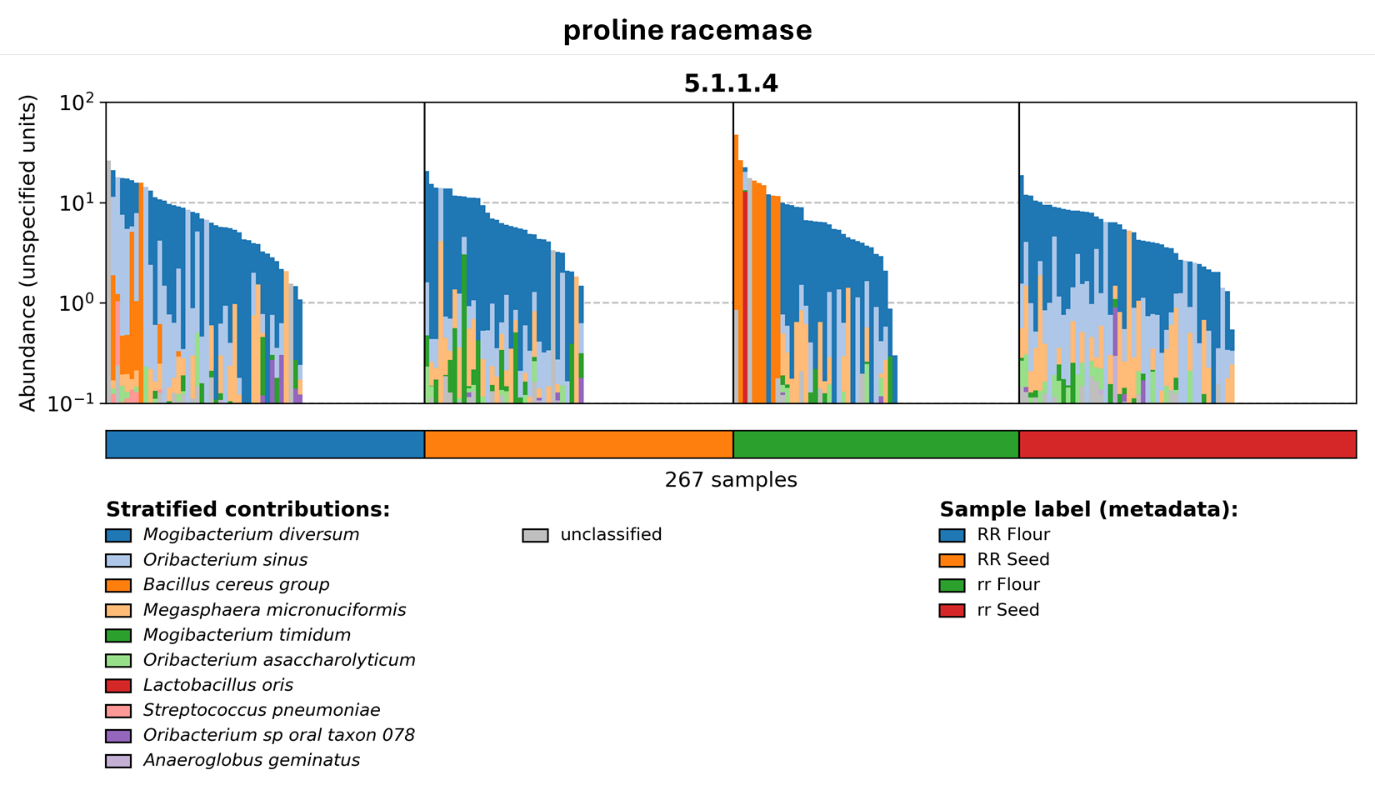

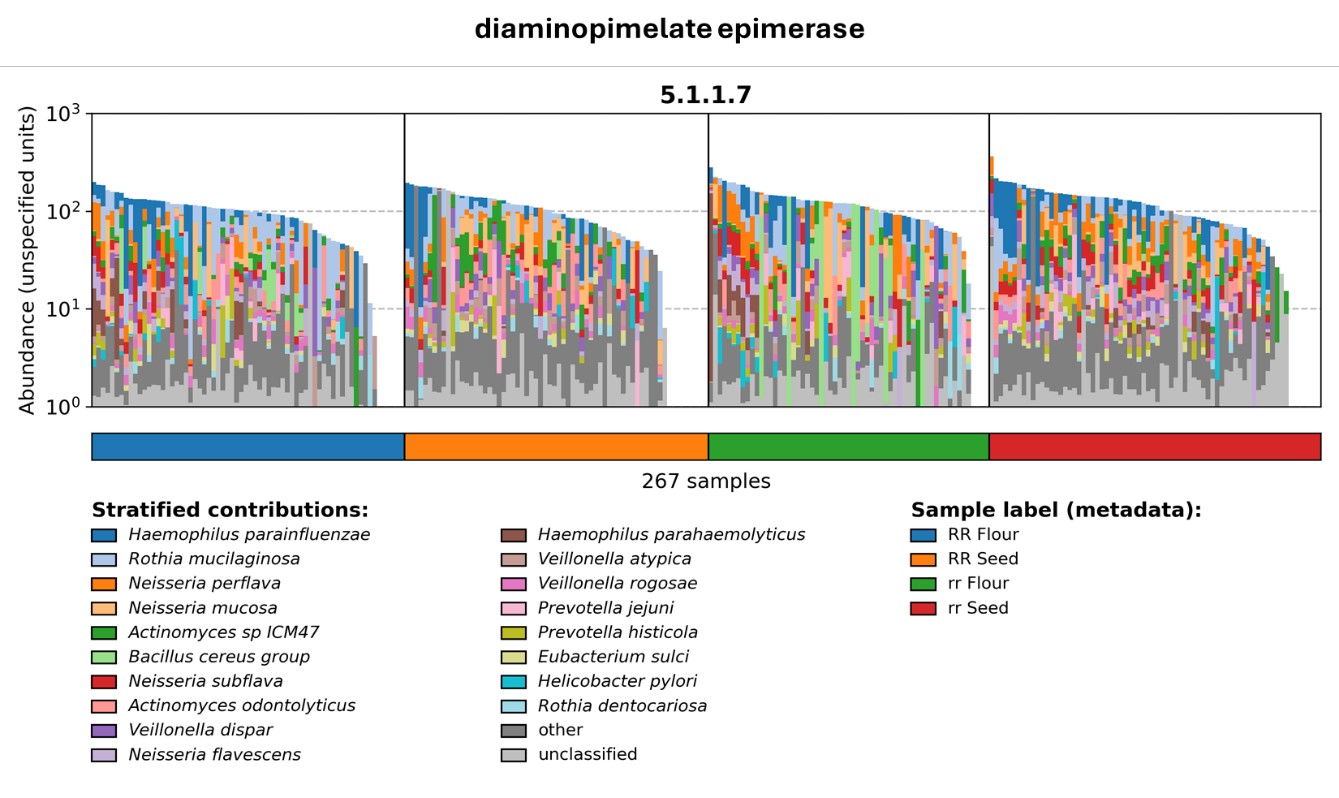

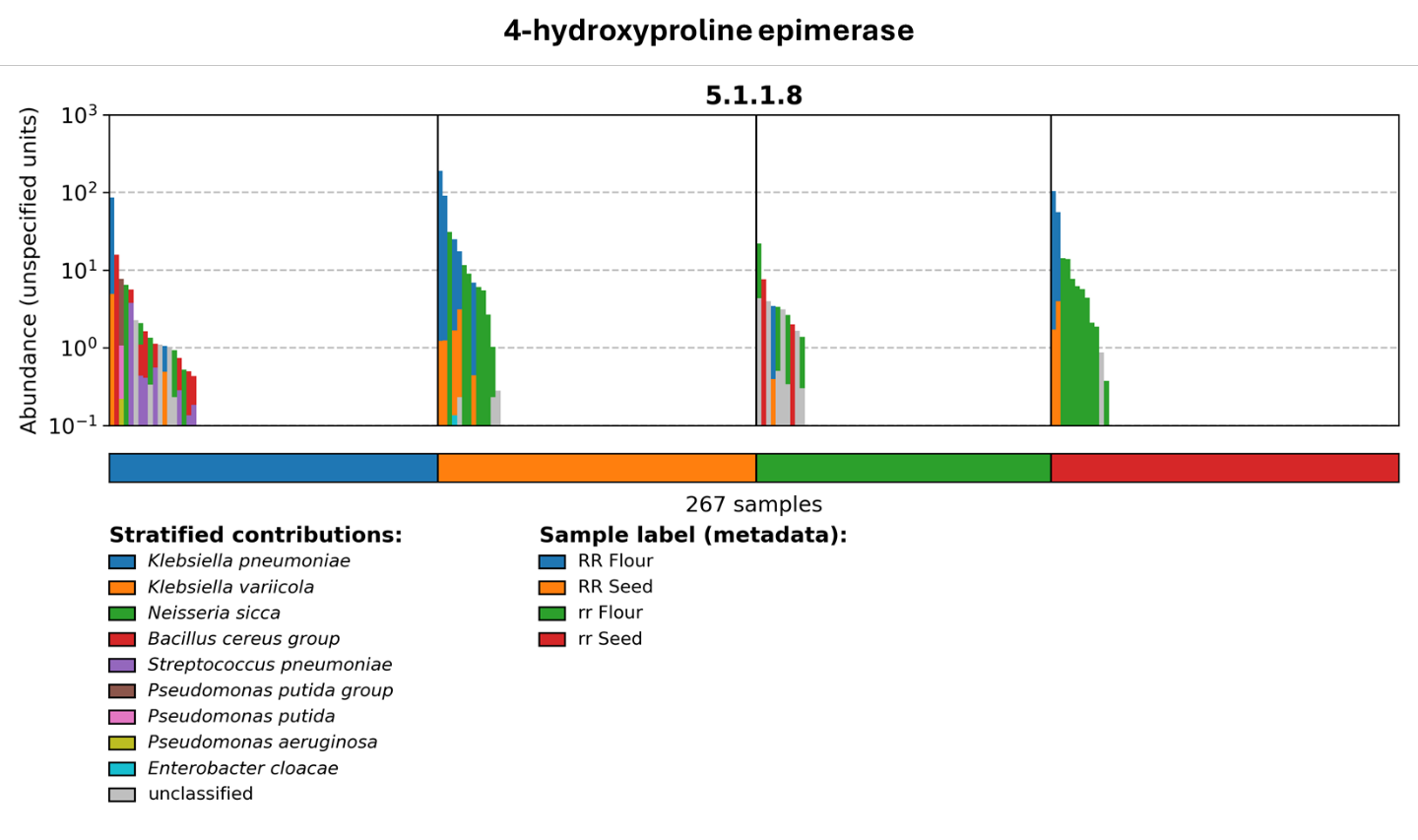

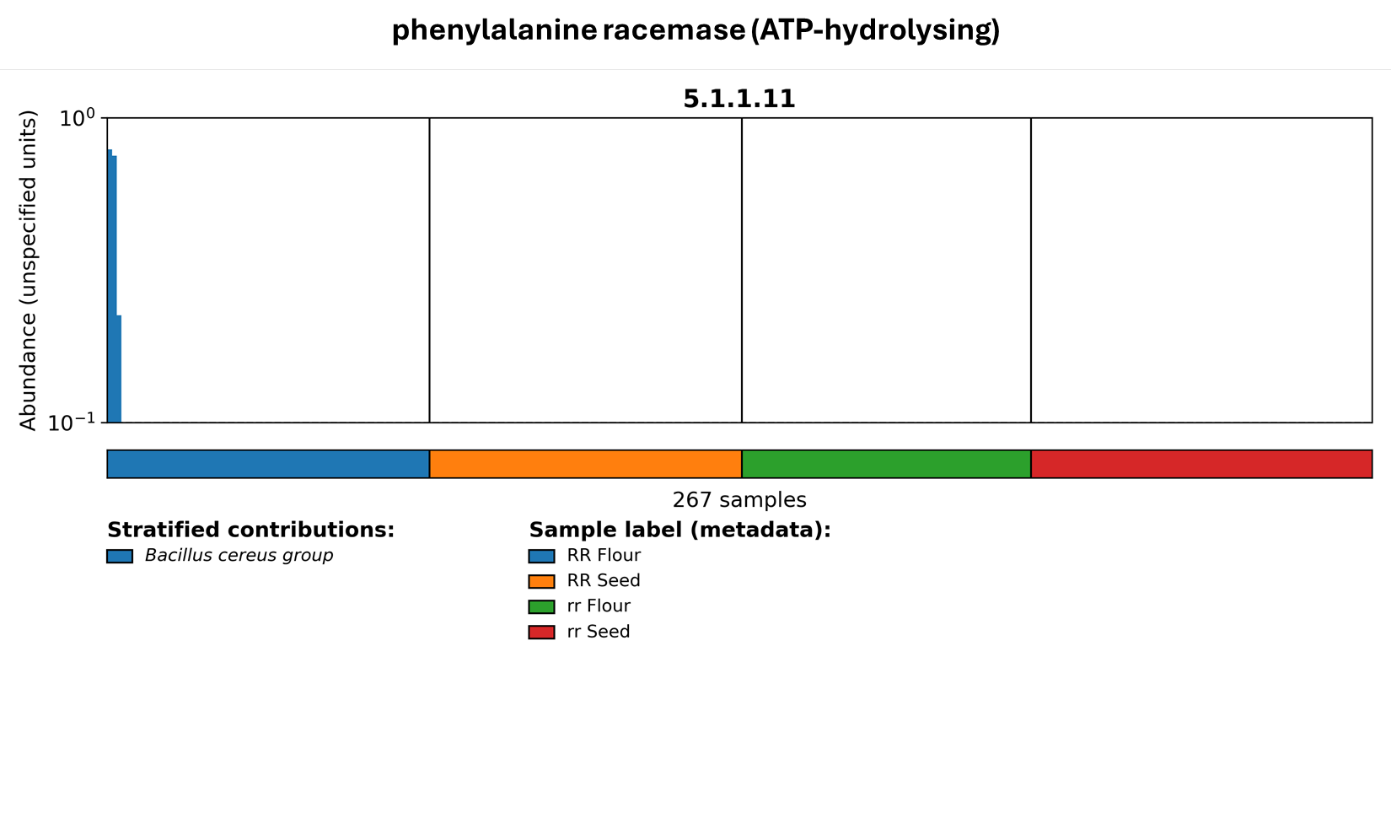

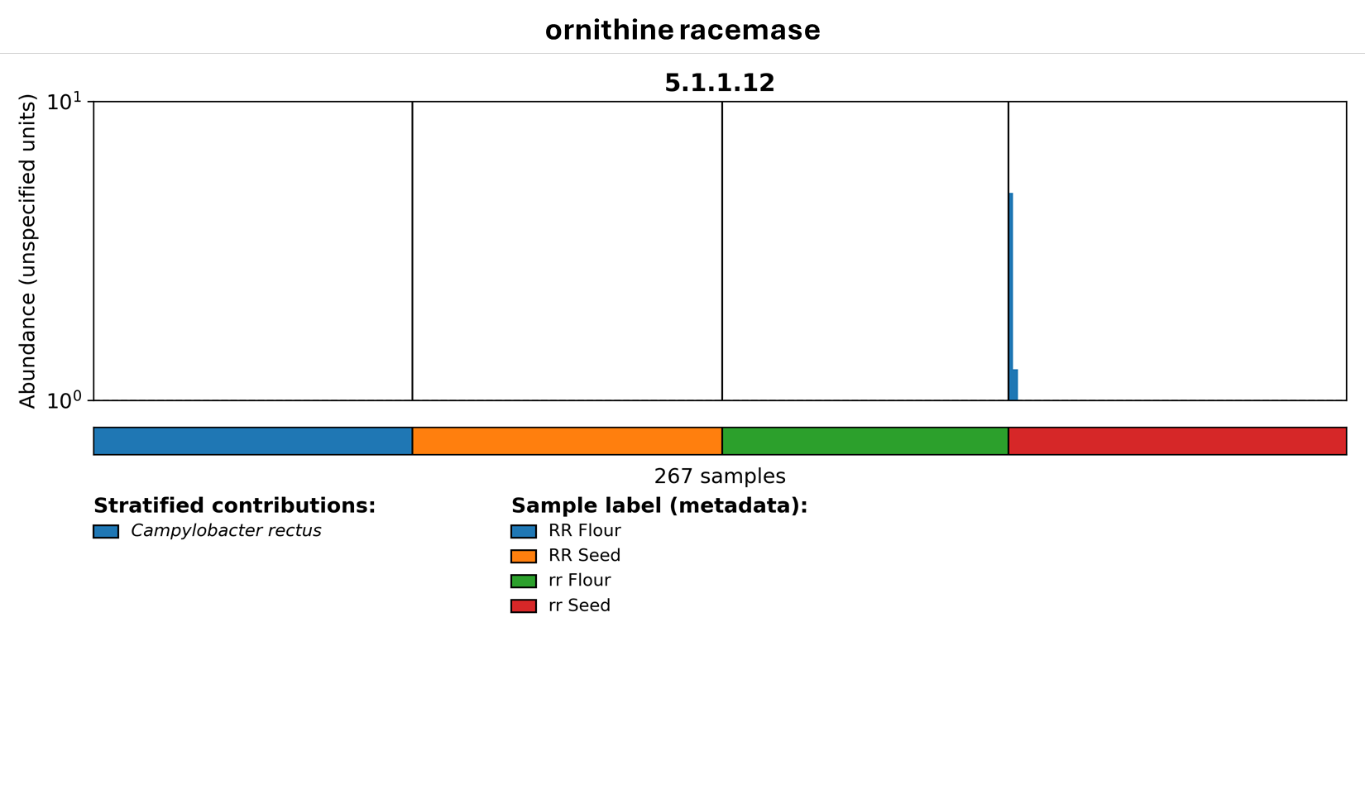

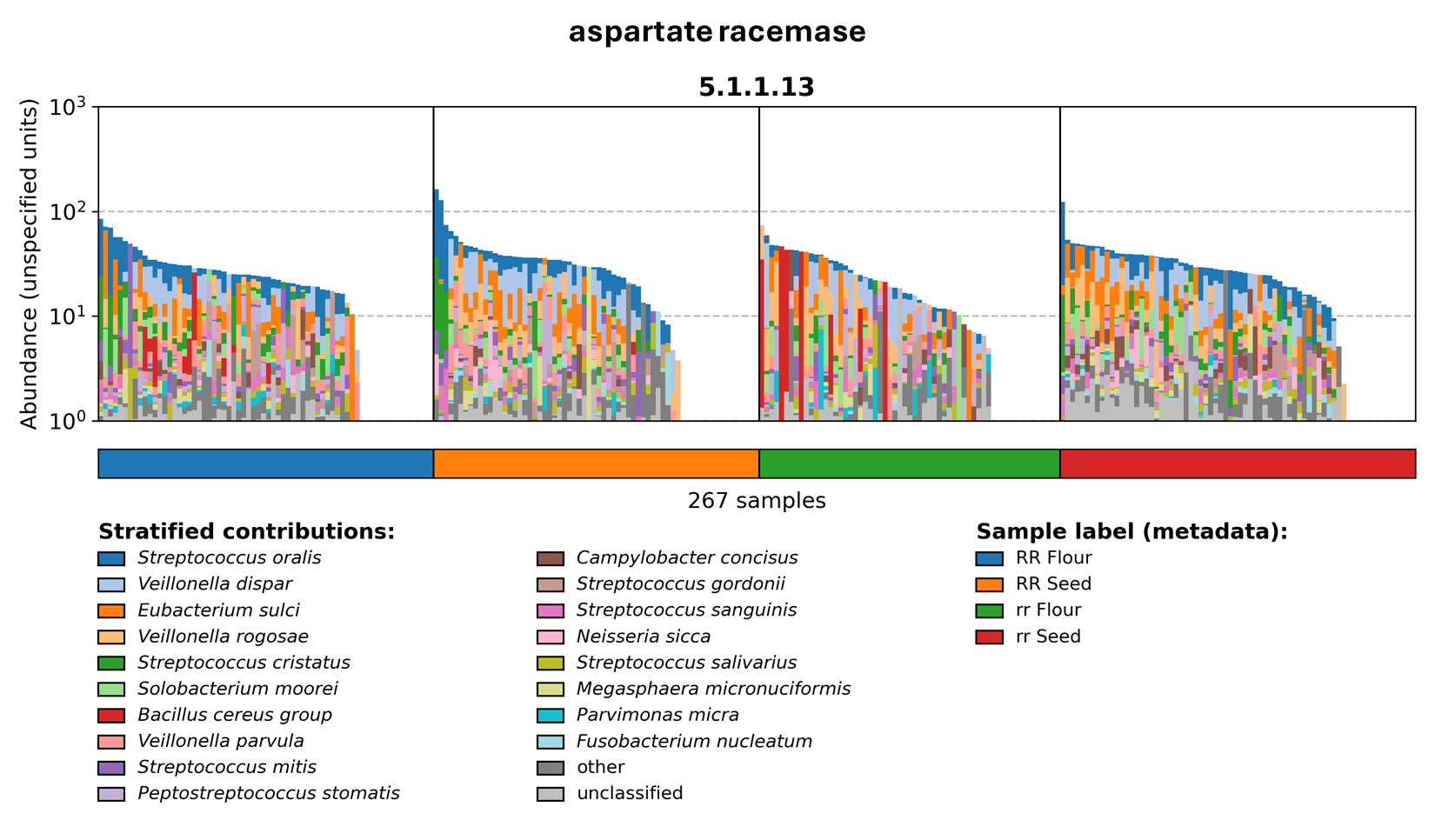

**Figure S6: Taxonomically stratified amino acid racemase gene abundances in the duodenal microbiome.** Amino acid racemase enzymes identified in stratified HUMAnN data through EC codes beginning with 5.1.1.X. The abundances are shown by meal, and are coloured by contribution from individual taxa.

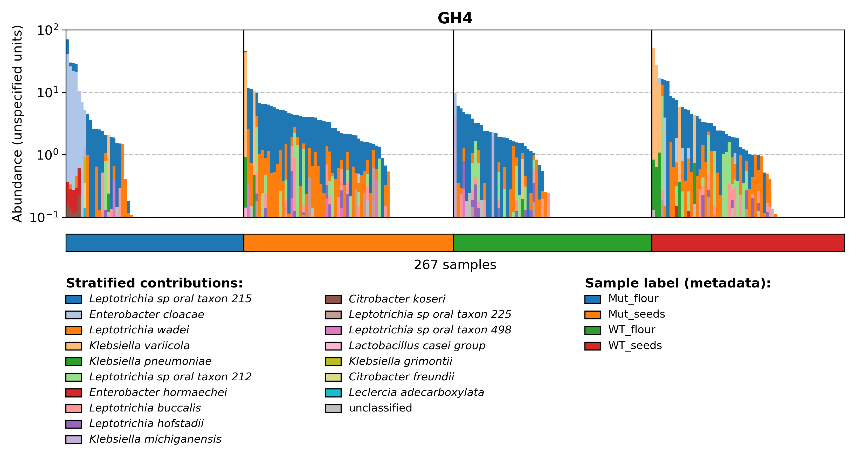

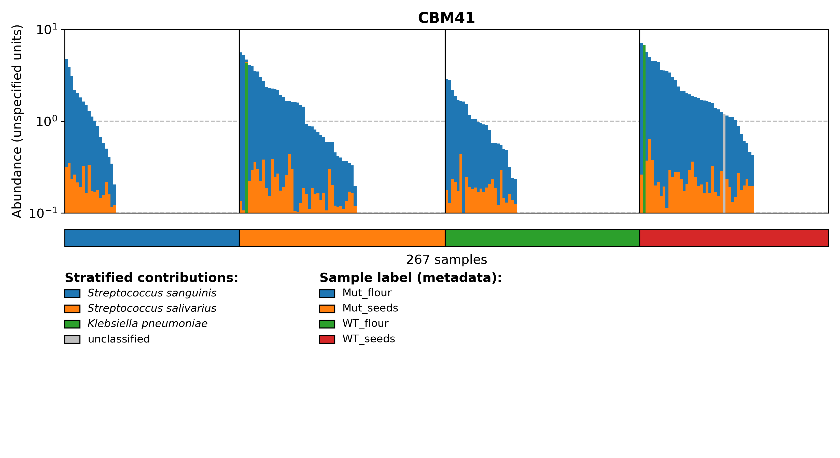

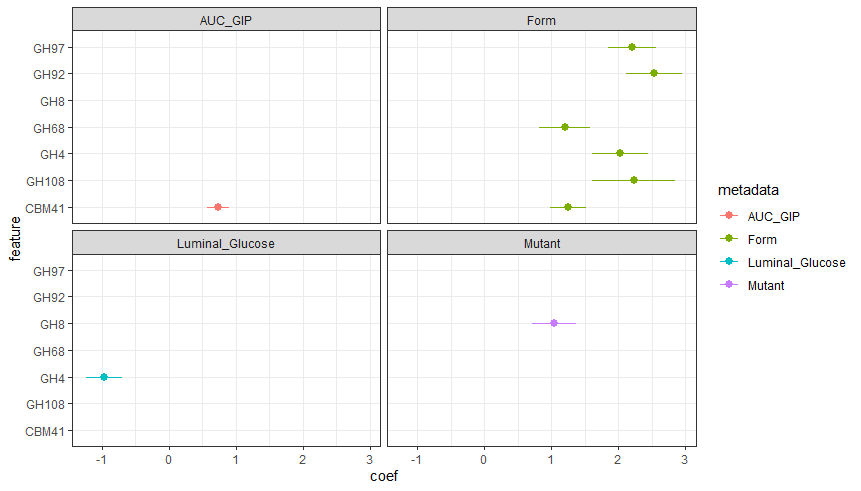

**Figure S7. Associations between duodenal microbiome CAZyme profiles,** **gut hormones, pea meal form and luminal glucose concentrations**. **A.** Statistically significant associations between gut microbial species (q < 0.05) and the post-prandial AUC for the hormone GIP; pea meal physical form (where a positive association demonstrates higher abundance in whole pea seed meals); luminal glucose concentrations in the duodenum; and pea genotype (where a positive association demonstrates higher abundance in the RR seed). **B.** CAZyme abundances for CBM41 and GH4 in each of the meals, stratified by the contribution of individual species.
